# Integrated single-nuclei and spatial transcriptomic analysis reveals propagation of early acute vein harvest and distension injury signaling pathways following arterial implantation

**DOI:** 10.1101/2023.10.31.564995

**Authors:** Marina E. Michaud, Lucas Mota, Mojtaba Bakhtiari, Beena E. Thomas, John Tomeo, William Pilcher, Mauricio Contreras, Christiane Ferran, Swati Bhasin, Leena Pradhan-Nabzdyk, Frank W. LoGerfo, Patric Liang, Manoj K. Bhasin

## Abstract

**Background:** Vein graft failure (VGF) following cardiovascular bypass surgery results in significant patient morbidity and cost to the healthcare system. Vein graft injury can occur during autogenous vein harvest and preparation, as well as after implantation into the arterial system, leading to the development of intimal hyperplasia, vein graft stenosis, and, ultimately, bypass graft failure. While previous studies have identified maladaptive pathways that occur shortly after implantation, the specific signaling pathways that occur during vein graft preparation are not well defined and may result in a cumulative impact on VGF. We, therefore, aimed to elucidate the response of the vein conduit wall during harvest and following implantation, probing the key maladaptive pathways driving graft failure with the overarching goal of identifying therapeutic targets for biologic intervention to minimize these natural responses to surgical vein graft injury.

**Methods:** Employing a novel approach to investigating vascular pathologies, we harnessed both single-nuclei RNA-sequencing (snRNA-seq) and spatial transcriptomics (ST) analyses to profile the genomic effects of vein grafts after harvest and distension, then compared these findings to vein grafts obtained 24 hours after carotid-cartoid vein bypass implantation in a canine model (n=4).

**Results:** Spatial transcriptomic analysis of canine cephalic vein after initial conduit harvest and distention revealed significant enrichment of pathways (*P* < 0.05) involved in the activation of endothelial cells (ECs), fibroblasts (FBs), and vascular smooth muscle cells (VSMCs), namely pathways responsible for cellular proliferation and migration and platelet activation across the intimal and medial layers, cytokine signaling within the adventitial layer, and extracellular matrix (ECM) remodeling throughout the vein wall. Subsequent snRNA-seq analysis supported these findings and further unveiled distinct EC and FB subpopulations with significant upregulation (*P* < 0.00001) of markers related to endothelial injury response and cellular activation of ECs, FBs, and VSMCs. Similarly, in vein grafts obtained 24 hours after arterial bypass, there was an increase in myeloid cell, protomyofibroblast, injury-response EC, and mesenchymal-transitioning EC subpopulations with a concomitant decrease in homeostatic ECs and fibroblasts. Among these markers were genes previously implicated in vein graft injury, including *VCAN* (versican), *FBN1* (fibrillin-1), and *VEGFC* (vascular endothelial growth factor C), in addition to novel genes of interest such as *GLIS3* (GLIS family zinc finger 3) and *EPHA3* (ephrin-A3). These genes were further noted to be driving the expression of genes implicated in vascular remodeling and graft failure, such as *IL-6*, *TGFBR1*, *SMAD4*, and *ADAMTS9.* By integrating the ST and snRNA-seq datasets, we highlighted the spatial architecture of the vein graft following distension, wherein activated and mesenchymal-transitioning ECs, myeloid cells, and FBs were notably enriched in the intima and media of distended veins. Lastly, intercellular communication network analysis unveiled the critical roles of activated ECs, mesenchymal transitioning ECs, protomyofibroblasts, and VSMCs in upregulating signaling pathways associated with cellular proliferation (MDK, PDGF, VEGF), transdifferentiation (Notch), migration (ephrin, semaphorin), ECM remodeling (collagen, laminin, fibronectin), and inflammation (thrombospondin), following distension.

**Conclusions:** Vein conduit harvest and distension elicit a prompt genomic response facilitated by distinct cellular subpopulations heterogeneously distributed throughout the vein wall. This response was found to be further exacerbated following vein graft implantation, resulting in a cascade of maladaptive gene regulatory networks. Together, these results suggest that distension initiates the upregulation of pathological pathways that may ultimately contribute to bypass graft failure and presents potential early targets warranting investigation for targeted therapies. This work highlights the first applications of single-nuclei and spatial transcriptomic analyses to investigate venous pathologies, underscoring the utility of these methodologies and providing a foundation for future investigations.

## Introduction

Coronary artery and peripheral arterial disease affect 20.1 million and 6.5 million adults in the United States, respectively.^1,2^ Both disease processes can lead to life-threatening ischemia, resulting in the need for surgical bypass surgery to redirect blood flow around blocked vessels. Autogenous vein, namely the greater saphenous vein, is the most widely used conduit for coronary artery bypass grafting (CABG) and is the gold-standard conduit for peripheral arterial bypass surgery.^3,4^ However, the 1-year primary patency rate of coronary and peripheral vein grafts is as low as 60%.^5,6^ The biological mechanisms of early vein graft failure (VGF) remain poorly understood but have been classically attributed to a maladaptive vascular wall remodeling response and the development of intimal hyperplasia (IH). Previously, to understand the molecular mechanism of IH in vein grafts, we performed a temporal analysis of bypass graft smooth muscle and endothelial cells following graft implantation.^7^ The innovative network-based analysis depicted that unresolved inflammation within 12 hours of graft implantation results in inflammatory and immune responses, apoptosis, proliferation, and extracellular matrix (ECM) reorganization in both cell types, leading to graft failure. Vein graft (VG) stenoses can occur anywhere along the vein conduit, suggesting that important injury response pathways may occur even earlier than previously described, namely in the preparation of autogenous vein grafts for implantation in the arterial system, as the conduit is manipulated and subjected to several sources of injury. Vein graft harvest injury includes ischemia of the vein wall during dissection from surrounding tissues with loss of vasa vasorum function in the vessel, pressure distension that induces hoop stress of the smooth muscle and endothelial cells, and exposure to an electrolyte solution, all of which may contribute to the upregulation of maladaptive pathways that persist and propagate following implantation into the arterial system. This response could be further exacerbated with distention of smaller diameter or poorly compliant phlebosclerotic veins.^8^ Accordingly, we aimed to identify the timeline associated with the upregulation of maladaptive pathways from initial distension during graft harvesting to implantation to set the groundwork for developing gene-therapy-based strategies to prevent or treat early vein graft injury and improve overall long-term vein bypass patency.

To accomplish this, we leveraged advances in spatial and single-cell technologies to provide a unique insight into the heterogeneity of the vascular landscape. Through the integrated use of single-nuclei RNA-sequencing (snRNA-seq) and spatial transcriptomics (ST) analysis, we examined the genomic effects of vein graft harvest, distention, and implantation across the regions of the vein, revealing distinct regulatory programs and cellular subpopulations mediating the response to acute distension injury.

## Materials and Methods

### Data availability

The sequencing data that supports the findings of this study are available via the Gene Expression Omnibus database under accession number (pending).

### Animals

All animal studies were approved by the Institutional Animal Care and Use Committee (IACUC) at Beth Israel Lahey Health and conducted in accordance with the National Institute of Health (NIH) Guide for the Care and Use of Laboratory Animals. Canines were obtained from Marshall Bioresources (North Rose, NY 14516, US), a registered class A vendor and AAALAC-accredited canine breeder for preclinical research purposes.

### Canine Surgery

Bilateral upper limb cephalic vein harvest was performed on 25 kg male or female mongrel dogs (n=4). General anesthesia was established and maintained with an initial sodium pentothal injection and subsequent 1% isofluorane inhalation after orotracheal intubation. Vein harvest was performed in conventional fashion. The cephalic vein was identified and exposed along the anterior surface from the wrist to the upper forearm. The distal portion of the vein was then ligated and transected. The end of the transected vein was cannulated with a metal cannula and secured with a silk suture. The vein was then sequentially distended by injecting a saline solution (0.9% sodium chloride, papaverine hydrochloride 60 mg, and 43 mg of bivalirudin) while occluding the outflow. Intervening tributaries were identified and ligated with silk sutures. A total length of 15 cm of vein was harvested. Total vein distension and harvest time was approximately 30 minutes. The cephalic vein following distention, along with a separate segment of the undistended cephalic vein, which served as the experimental control, were excised. Samples were collected and immediately snap-frozen in cryovials using liquid nitrogen for single-nuclei analysis or embedded in O.C.T. blocks and flash-frozen in an isopentane and liquid nitrogen bath for spatial transcriptomics analysis. Systemic bivalirudin was then infused before the creation of the anastomoses. Bilateral end-to-side anastomoses were created on the vein with 7-0 Prolene sutures. The intervening segment of the carotid artery was then ligated, creating preferential flow through the graft. After hemostasis was achieved, the wounds were closed in three layers with 3-0 vicryl sutures. The patency of the graft was confirmed through direct palpation and ultrasound imaging. Early (24 hours) and late (30 days) time points were chosen, and harvest was performed with the dog under general anesthesia. Vein graft samples were then collected and processed as previously described for cryopreservation and OCT embedding.

### Immunohistochemistry

Formalin-fixed and paraffin-embedded tissue sections from each sample were cut to a 5-μm thickness and air-dried. The staining for GLIS Family Zinc Finger 3 (GLIS3), Versican (VCAN), and fibrillin 1 (FBN1) was performed using the Leica Bond RXM automated immunohistochemistry staining platform (Leica Biosystems). Before staining, the slides were baked for 30 minutes in a 60 °C oven. The slides were then labeled and loaded onto the Bond RXM. The slides were deparaffinized with the Bond Dewax Solution and rinsed with Leica Wash Buffer. Following deparaffination, the slides were heated to 100 °C, and antigens were retrieved for 20 minutes with ER2 (high pH) antigen retrieval buffer and then rinsed with Leica Wash Buffer. The peroxidase block was applied at room temperature for 5 minutes, and the sections were washed with three rinses of wash buffer. The diluted antibodies (GLIS3 at 1:40, VCAN at 1:200, and FBN1 at 1:500) were applied to the slides for 30 min at room temperature, followed by three rinses of wash buffer. Leica Bond anti-Rb HRP secondary was applied and incubated for 8 min, and the detection was completed in combination with the Leica Refine DAB kit, as per manufacturer recommendations. Slides were counterstained with hematoxylin for 5 min. Slides were then dehydrated, cover-slipped, and evaluated by light microscopy with scanned images of the slides. The slides were scanned on a Hamamatsu Nanozoomer HT 2.0 at 40x.

### Single-molecule RNA in situ hybridization

RNA in situ hybridization experiments were performed using the RNAscope® technology,^9^ which has been previously described.^9^ Paired double-Z oligonucleotide probes were designed against target RNA using custom software. The following probes were used: Cl-LMCD1-C1 (Part ID: 1317341-C1), Cl-ELN-C1 (Part ID: 1317351-C1), and Cl-FBN1-C1 (Part ID: 1317361-C1). The RNAscope Multiplex Fluorescent Detection Kit v2 (cat. no. 323110) (Advanced Cell Diagnostics) was used according to the manufacturer’s instructions. Slides of flash-frozen tissue samples of control and distended veins were prepared according to the manufacturer’s recommendations. Each sample was quality-controlled for RNA integrity with a probe specific to the housekeeping genes *Polr2a, PPIB,* and *UBC*. Negative control background staining was evaluated using a probe specific to the bacterial *dapB* gene. Fluorescent images were acquired using a Leica Stellaris X5 confocal microscope with a 40x objective.

### Single-nuclei transcriptomics sample preparation

We used the sample prep user guide (10x Genomics, CG000505, Rev A) for the Chromium Nuclei Isolation Kit with RNase Inhibitor (10x Genomics, 1000494) to isolate nuclei from flash-frozen tissues. The various buffers (lysis, debris removal, wash, and resuspension buffers) were prepared using kit components according to user guide instructions (10x Genomics, CG000505, Rev A). Briefly, ∼50 mg of tissue, cut into smaller pieces to enable better lysis of cells, was added to pre-chilled sample dissociation tubes and dissociated by adding lysis buffer followed by multiple strokes with the pestle against the dissociation tubes. Following incubation on ice for 10 minutes, the dissociated tissue was added to the nuclei isolation column, placed in a pre-chilled collection tube, and spun at 16,000 g for 20 sec at 4°C. The flowthrough was collected, vortexed to resuspend the nuclei, and spun at 500g, 3 min, 4°C. The pellet was then resuspended in debris removal buffer, pipet mixed, and spun at 700 g, 10 min, 4°C. After discarding the supernatant, the pellet was washed with wash and resuspension buffer by spinning at 500 g, 5min, 4°C. For counting, the nuclei were stained with acridine orange/propidium iodide stain (Logos Biosystems) and counted using a LUNA-FX7 cell counter (Logos Biosystems). The nuclei pellet was resuspended in wash and resuspension buffer to get the desired nuclei concentration (700-1200 nuclei/ml). The snRNA-seq libraries were prepared immediately from the isolated nuclei with the Chromium Next GEM Single Cell 3’ v3.1 reagent kits (10x Genomics, 1000268). Briefly, GEMs were prepared using the 10x Genomics controller followed by RT-PCR and cDNA amplification. The cDNA was then used to prepare snRNA-seq libraries according to the user guide (10x Genomics, CG000315 rev. B). The quality of the cDNA and final library preps was assessed using DNA HS bioanalyzer kits (Agilent), and concentration was estimated using a Qubit fluorometer (Thermo Fisher Scientific). Sequencing was performed using the massively parallel sequencing on the Novaseq 6000 platform using Novaseq S4 PE 100 kits (Illumina) to capture the expression of ∼200-5,000 genes/nuclei.

### Visium tissue permeabilization optimization, gene expression library construction, and sequencing

Frozen samples embedded in O.C.T. (TissueTek Sakura) were cryosectioned (10 μm sections) at −20 ^°^C and transferred onto pre-chilled Visium Tissue Optimization Slides (10x Genomics, 3000394) and Visium Spatial Gene Expression Slides (10x Genomics, 2000233). H&E staining was performed according to the Visium Spatial Gene Expression User Guide (10x Genomics, CG000239 rev. F). Optimal permeabilization time was determined using the tissue optimization slide, which was processed according to the optimization protocol (10x Genomics, CG000239 rev. F). The tissue was permeabilized with permeabilization enzyme for different periods of time (6, 9, 12, 18, 24, and 30 minutes). Brightfield H&E images were taken using a 40X objective on Nanozoomer 2.0 HT. For tissue permeabilization optimization experiments, fluorescent images were taken using a 20X objective (0.5 µm/pixel) and CY3 and CY5 MSI filters at 177.75 ms and 73.31 ms exposure time in an Akoya Vectra Polaris whole slide scanner. The permeabilization time with maximum fluorescent signal was taken as the optimal time for permeabilization when processing gene expression slides. Brightfield H&E images were taken using a 40X objective on Nanozoomer 2.0 HT. For tissue permeabilization optimization experiments, fluorescent images were taken using a 20X objective (0.5 µm/pixel) and CY3 and CY5 MSI filters at 177.75 ms and 73.31 ms exposure time in an Akoya Vectra Polaris whole slide scanner. For the Visium spatial transcriptomics gene expression assay, tissue was permeabilized for the optimal time (9 minutes) followed by reverse transcription, second strand synthesis and denaturation, cDNA amplification and QC, and gene expression library construction according to 10x Genomics user guide (CG000239 rev. F). The gene expression libraries were quantified using a Qubit fluorometer (Thermo Fisher Scientific) and checked for quality using DNA HS bioanalyzer chips (Agilent). Sequencing depth was calculated based on the percent capture area covered by the tissue, and the 10x Genomics recommended sequencing parameters were used to run on the Novaseq S4 PE 100 kits (Illumina).

### Transcriptional profiling and data analysis

FASTQ files for each sample were aligned to the Dog10K_Boxer_Tasha_1.0 canine reference genome^10^ using Space Ranger (v2.0.0, 10x Genomics) for spatial transcriptomics samples (n=7) and Cell Ranger (v7.0.0, 10x Genomics) for single-nuclei samples (n=4). For spatial samples, microscopy images of the H&E stained samples were aligned to the spatial voxels through the Space Ranger pipeline. The filtered feature-barcode matrices for the single-nuclei samples were imported into Seurat (v4.3.0),^11^ then low-quality nuclei with <200 unique genes, <500 UMI reads, or ≥30% mitochondrial transcripts were filtered out. Due to the increased number of nuclei retrieved from the distended vein of Dog 2, this sample was downsampled to 1,000 nuclei that were used for subsequent analysis, keeping both the control vein and distended vein groups similar in total nuclei (1,233 and 1,188 nuclei, respectively) for comparison. For both the single-nuclei and spatial datasets, the samples were normalized with SCTransform and then integrated using integration anchors-based batch correction via Seurat. Principal component analysis (PCA) was then performed on the integrated spatial and single-nuclei datasets using all non-ribosomal features, followed by construction of a K-nearest neighbors graph (using cosine similarity) and Louvain clustering, then non-linear dimensional reduction via Uniform Manifold Approximation and Projection (UMAP). The resolution and number of principal components used for clustering were selected using the clustree R package (v0.5.0).^12^ Differential gene expression analysis was performed via Seurat using the FindAllMarkers function, comparing expression between clusters or sample groups (distended veins versus control veins).

### Statistical analysis and data visualization

All statistical analysis was performed using R. Significantly differentially expressed genes were determined based on the Wilcoxon rank test with Bonferroni multiple test correction (*P* < 0.05). Significantly enriched pathways were calculated using Benjamini-Hochberg adjusted p-values (*P adj (BH)* < 0.05). Spatial plots, UMAP plots, and differential gene expression heatmaps/dot plots were all generated using Seurat. Bar plots of the cluster distributions and the deconvolution matrix were generated using ggplot2 (v3.4.1).^13^ Volcano plots based on differential gene expression were made using the EnhancedVolcano R package (v3.16). Dot plots of enriched pathways were generated via the dotplot function in clusterProfiler. Networks of enriched pathways were generated via the cnetplot function in the clusterProfiler Bioconductor package.^14^

## Results

### Optimization of tissue permeabilization techniques for the preparation of vein samples for spatial and single-nuclei transcriptomic analyses

Bilateral upper limb cephalic vein harvest was performed on canine specimens to yield three matched pairs of control and distended veins (Figure 1A). Following distension of the cephalic vein with a saline solution containing papaverine and bivalirudin, segments of the vein were excised along with separate undistended cephalic vein segments. These tissue samples were subsequently cryopreserved in liquid nitrogen for single-nuclei analysis or embedded in O.C.T. and flash-frozen for spatial transcriptomic assays. Single-nuclei methodologies uniquely enable the transcriptomic analysis of challenging vascular tissues by eliminating the need to maintain whole-cell integrity during tissue dissociation while still preserving individual cellular resolution at the transcriptomic level. Therefore, to overcome the challenges associated with obtaining viable single-cell suspensions from fibrous vascular tissues, ^15^ we endeavored to harness snRNA-seq to analyze the vein sample transcriptomes. Through this approach, we were successfully able to capture diverse cell types within the vein (i.e., smooth muscle cells, fibroblasts, endothelial cells, and leukocytes) with 200-1,000 nuclei expressing ∼200-5,000 genes per sample, illustrating single-nuclei isolation as a viable alternative to single-cell isolation for the transcriptomic analysis of difficult-to-process vascular tissue (Table S1).

**Figure 1.**
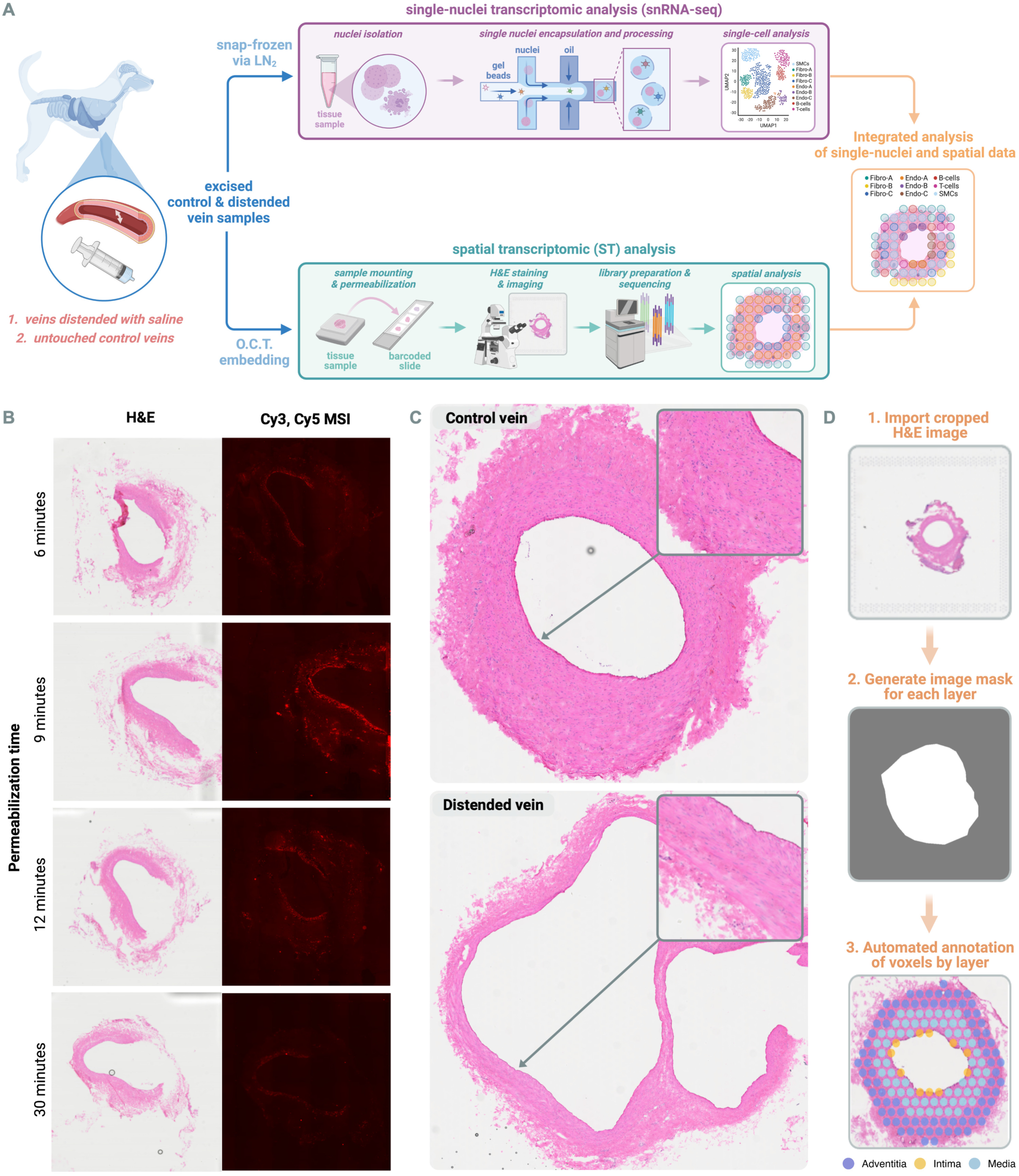
Overview of the workflow used for sample analysis. **(A)** Canine vein graft samples were retrieved following distension, then analyzed via single-nuclei RNA-seq (snRNA-seq) and spatial transcriptomics (ST) through the 10x Genomics platform. Single-nuclei-based transcriptomics enables the sequencing of difficult-to-dissociate samples while preserving detailed individual cellular resolution. As a complementary approach, ST provides insight into the spatial gene expression of the tissue, with the transcriptomes of 1-10 cells captured per barcoded spot of the sample slide. Combined, these approaches illuminate the unique cellular architecture and transcriptomic landscape of the tissue. **(B) Varying permeabilization times were assessed for fluorescent cDNA footprint to optimize tissue permeabilization**. The left panel shows the brightfield images (H&E), and the right panel shows fluorescent images (Cy3, Cy5 Multispectral Imaging (MSI)). **(C)** Images of H&E strained veins. Notably, the lower panel illustrates the expansion of the lumen following distension. **(D)** Overview of the developed method for adding histological annotations to ST samples for targeted downstream analyses. Capture spots are projected onto the H&E image as voxels, generating a spatial plot that can be annotated with STannotate and used for analysis.

As this work presents the first spatial transcriptomic investigations of superficial vein samples, we first sought to optimize the tissue permeabilization time for spatial transcriptome profiling. Using the Visium platform from 10x Genomics, tissue sections are transferred and fixed onto capture areas containing ∼5,000 barcoded spots; cDNA synthesis subsequently incorporates these barcodes to ultimately facilitate the transcriptomic analysis of tissue by spot (1-10 cells captured per spot) (Figure 1A). Tissue permeabilization presents a crucial step in the sample preparation, as insufficient permeabilization will reduce the amount of RNA released and captured, thus reducing the transcriptomic resolution and affecting downstream analyses, while over-permeabilization results in cDNA diffusion, compromising the spatial localization.^16^ Examining six different permeabilization times (6–30 minutes) via fluorescence microscopy (Figure 1B), we identified 9 minutes as the optimal permeabilization time for 10 µM vein tissue based on the fluorescent signal. Applying this technique, we successfully performed spatial transcriptomics on the vein samples, yielding 125-550 spots captured per sample with an average expression of ∼870 genes per spot (Table S1).

Lastly, to facilitate targeted spatial transcriptomic analysis of the veins, we developed a facile computational method for annotating the spatial plot voxels according to histological analysis (Figure 1C). To accomplish this, we utilized the open-source imaging software ImageJ to produce image masks for each layer of the vein wall, then developed an R-based function for adding these labels to the metadata of each spatial sample (Figure 1D). This function, which will be available through the STannotate package, enables researchers to readily add histological annotations to vein samples for downstream analysis.

### Integration of single-nuclei and spatial transcriptomics illustrates a significant impact of distension on enrichment and profiles of endothelial and fibroblast subpopulations

To investigate how distension alters the spatial gene expression landscape of the vein, we first performed spatial transcriptomic (ST) analysis on cephalic vein samples from four canines, including three vein samples retrieved following distension with saline and four untouched control vein samples (Table S1). Applying our STannotate tool, we labeled the spatial voxels of each sample into intimal, medial, and adventitial layers based on the tissue morphology observed through histological analysis (Figure S1). After this, we performed a comparative analysis of the transcriptomic profiles of the intimal, medial, and adventitial layers between distended and control veins (Figure 2A).

**Figure 2.**
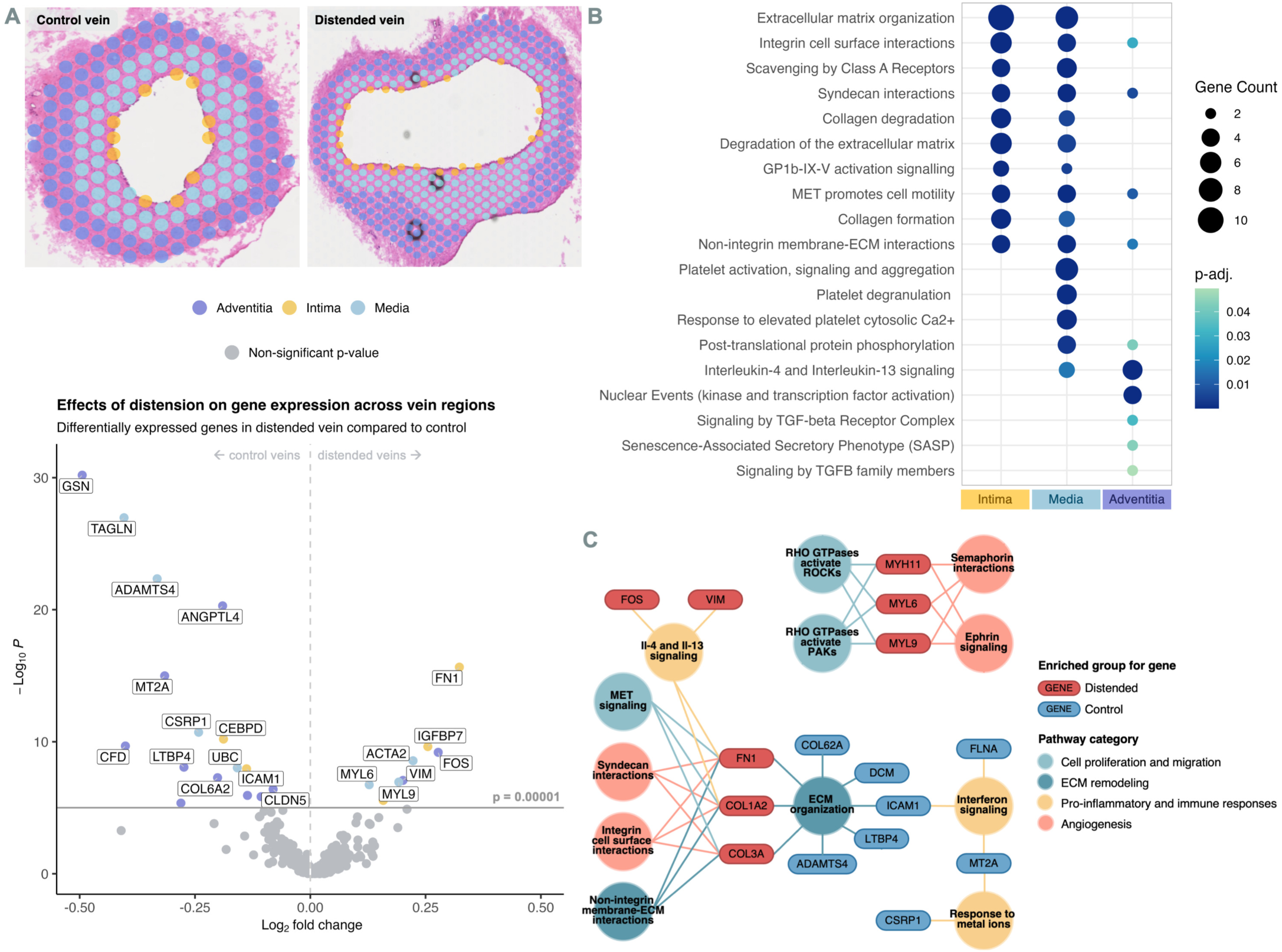
Spatial transcriptomic analysis of distended veins. **(A)** Top: Spatial plots displaying capture spots (voxels) projected onto H&E images of control (left) and distended (right) veins from the same canine, annotated using our ST annotation tool. Bottom: Volcano plot of significantly differentially expressed genes (*P* < 0.00001) within the distended vein compared to the control vein, wherein each gene is colored according to the vein wall layer with the highest expression of the marker. **(B)** Enriched pathways (via Reactome database) within each layer of the distended veins compared to control veins, based on significantly upregulated genes between each layer (*P* < 0.05). **(C)** Network of key differentially expressed genes and associated pathways (*P* < 0.05) between control and distended veins, derived from clusterProfiler analysis using the Reactome pathways database.

Following distension, significant overexpression of genes involved in extracellular matrix (ECM)-remodeling, cell proliferation, and migration, as well as cytokine production, were found to be overall significantly upregulated (*P* < 0.00001) in the vein conduit. Specifically, within the intima, regulators of endothelial cell (EC) proliferation and migration, fibronectin (*FN1*) and insulin-like growth factor-binding protein 7 (*IGFPB7*),^17,18^ were transcriptionally upregulated. Similarly, in the adventitia, genes associated with fibroblast (FB) activation and cytokine production, c-Fos (*FOS*) and vimentin (*VIM*), were upregulated. Within the media, transcription of the contractile VSMC markers, α-SMA (*ACTA2*) and myosin light chain proteins (*MYL6* and *MYL9*), were increased. Expression of contractile VSMC markers is typically associated with a quiescent SMC phenotype; however, a significant decrease in the contractile markers transgelin (*TAGLN*) and cysteine-rich protein 1 (*CSRP1*) was concurrently observed, conversely signifying VSMC-preprogramming to a synthetic (activated) phenotype (Figure 2A, bottom panel). Furthermore, among the downregulated genes in the distended veins, negative regulators of oxidative stress, metallothionein 2A (*MT2A*) and angiopoietin-like 4 (*ANGPTL4*), were decreased in the adventitia, suggesting that oxidative stress response may be dysregulated in distended veins (Figure 2A, bottom panel).

We next examined the pathways enriched within each layer of the distended vein by performing pathway enrichment analysis (Figure 2B) on the differentially expressed genes (*P* < 0.05) and mapping the network of enriched pathways between distended and control veins (Figure 2C). The results of this pathway analysis further illustrate significantly increased (*P* < 0.05) ECM remodeling, cell proliferation and migration, and cytokine production following distension, wherein integrin- and syndecan-mediated interactions and collagen degradation and formation, as well as increased Rho GTPase-, ephrin-, and MET-mediated cell motility, were elevated in the intima and media. Additionally, the media displayed significant enrichment of platelet activation and profibrotic IL-4 and IL-13 cytokine signaling (*P* < 0.05),^19^ the latter of which was also enriched in the adventitia (Figure 2B).

Collectively, these initial results of our ST analysis suggest that distension of the vein initiates the upregulation of pathways implicated in the activation of ECs, VSMCs, and FBs, namely ECM remodeling across the inner vein wall, platelet activation driven by the media, cytokine signaling within the outer vein wall, and the increased migratory signaling across all three layers.

#### Clustering of spatial voxels reveals the heterogeneous distribution of transcriptional profiles driving cellular activation and vascular remodeling within the media and adventitia following distension

To expand upon these findings and investigate the spatial transcriptomic landscape through an unbiased approach, we next clustered the spatial voxels (containing 1-10 cells each) based on their cell-type composition and transcriptomic profiles via K-nearest neighbor (KNN) graph construction and Louvain clustering, visualized by Uniform Manifold Approximation and Projection (UMAP) (Figure 3A). Seven clusters formed, which we subsequently labeled according to their primary localization to the intima (e.g., I1), media (e.g., M1), or adventitia (e.g., A1). Markedly, the spatial distribution of these clusters within the distended vein (Figure 3B, Figure S2) was nonuniform compared to the control vein (Figure 3C, Figure S2), highlighting the heterogeneous effect of sheer stress during distension that may contribute to dysregulated healing in segments of the vein, leading to IH progression observed in graft failure.^20^

**Figure 3.**
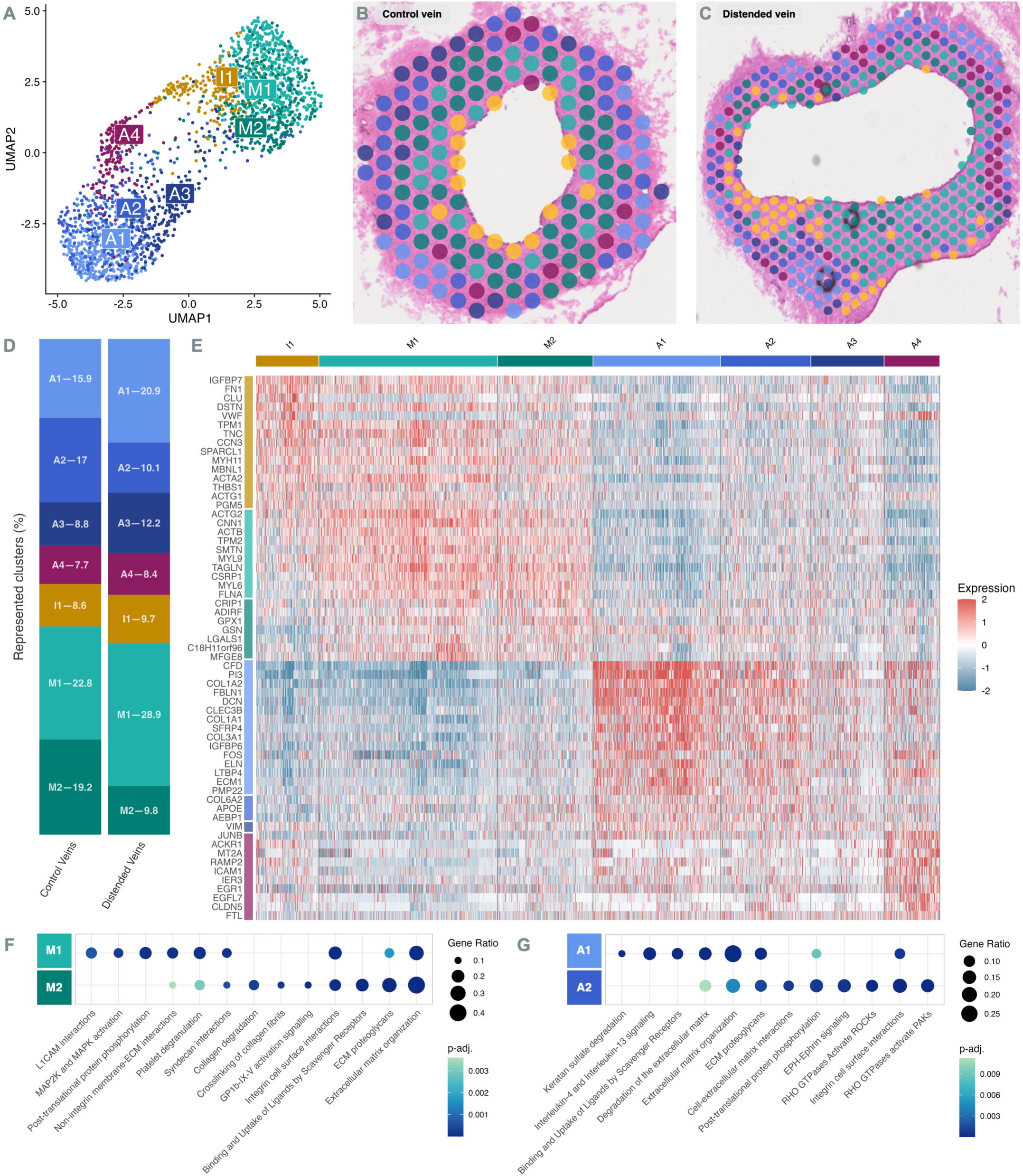
Spatial clustering of control and distended veins. **(A)** UMAP displaying the clustering of spatial transcriptomic voxels (containing 1-10 cells each) derived from control and distended veins. Clusters are derived based on similar transcriptomic profiles correlating to similar cell type compositions. Cluster labels were designated by the predominant localization of associated voxels to either the intima (I), media (M), or adventitia (A) of the vein samples. The spatial distribution of these clusters was then mapped onto the vein samples. **(B)** Mapping of spatial clusters onto a control vein sample. **(C)** Mapping of spatial clusters onto a distended vein sample from the same canine. **(D)** Relative proportions of each spatial cluster present within control vein and distended vein groups. (E) Heatmap of marker genes associated with different spatial clusters. Columns represent individual voxels grouped by spatial cluster, while rows display individual genes. Horizontal colored bars above the heatmap indicate the different spatial cellular clusters. Relative gene expression is shown in pseudo color, where blue represents low expression and red represents high expression. Enriched biological pathways (via clusterProfiler using Reactome database) based on differentially upregulated genes between the (**F**) M1 and M2 clusters, as well as the **(G)** A1 and A2 clusters (*P adjusted <* 0.05). Gene ratio is defined as the proportion of upregulated genes present in the cluster that overlap with the respective pathway. The multiple test correction for the P value has been performed using the Benjamini-Hochberg (BH) approach.

Particularly, while the intimal cluster (I1) evenly surrounded the lumen (Figure 3B) in the control vein, this cluster significantly extended into portions of the media following distension (Figure 3C). Notably, this cluster depicted significant upregulation (*P* < 0.00001) of EC activation and endothelial-mesenchymal transition (EndMT) associated genes, namely fibronectin (*FN1*),^21^ von Willebrand factor (*VWF*),^22^ tropomyosin 1 (*TPM1*),^23^ α-SMA (*ACTA2*),^24^ γ-actin (*ACTG1*),^25^ and thrombospondin 1 (*THBS1*),^26^ suggesting that distension promotes the emergence of activated EC and mesenchymal phenotypes (Figure 3E).

Furthermore, distended veins became enriched in their proportion of clusters A1 and M1 and correspondingly decreased in clusters A2 and M2 (Figure 3D), indicating a shift in genomic expression within the media and adventitia following distension. While clusters M1 and M2 shared similar expression profiles, the M1 cluster (enriched in distended veins) displayed modest upregulation (*P* < 0.00001) of contractile VSMC marker genes (Figure 3E). Further pathway enrichment analysis showed significant enrichment (*P adj (BH)* < 0.005) of cell activation and migration pathways, including ECM remodeling, MAPK signaling, L1CAM-mediated cell adhesion, and platelet degranulation (Figure 3F). In contrast, the M2 cluster (enriched in control veins) primarily showed significant upregulation (*P* < 0.00001) of genes negatively regulating proliferation, such as cysteine-rich protein (*CRIP1*)^27^, antioxidant glutathione peroxidase-1 (*GPX1*),^28^ adipogenesis regulatory factor (*ADIRF*), and gelsolin (*GSN*) (Figure 3E). Notably, gelsolin was found to be the most significantly downregulated gene (P < 0.00001) overall in distended veins (Figure 2A). While previous work has demonstrated that gelsolin inhibits angiotensin II-mediated activation of myofibroblasts in mycardiofibrosis,^29^ its role in vascular injury has not been elucidated and warrants future investigation. Within the adventitia, the A1 cluster displayed significant upregulation (*P* < 0.00001) for key regulators of ECM remodeling, including collagen I (*COL1A*) and collagen III (*COL3A*), fibulin-1 (*FBLN*1), elastin (*ELN*), and extracellular matrix protein 1 (*ECM1*) (Figure 3E). In contrast, the A2 cluster showed significant over-expression (*P* < 0.00001) of genes implicated in fibroblast-mediated healing, including negative regulator of cell migration apolipoprotein E (*APOE*),^30^ profibrotic collagen VI (*COL6A2*),^31^ and vascular-remodeling regulator aortic carboxypeptidase-like protein (*ACLP/AEBP1*) (Figure 3E).^32^ Furthermore, pathway enrichment analysis illustrated significantly increased activation (*P adj (BH)* < 0.01) of ECM remodeling and cytokine signaling pathways in the A1 cluster, while the A2 cluster exhibited significant activation (*P adj (BH)* < 0.01) of ROCKs, PAKs, and ephrin kinases involved in modulating cell migration, potentially aiding in fibroblast-mediated healing (Figure 3G).

Together, these results of spatial clustering, coupled with the initial ST analysis, demonstrate that distension alters the transcriptomic landscape of the vein, coordinating a distinct response to the imparted endothelial injury across each layer of the vein wall. Specifically, within the intima and media, markers of EC activation, cellular proliferation and migration, platelet activation, and EndMT are primarily upregulated following distension, while in the adventitia, key markers of fibroblast-mediated ECM remodeling are upregulated.

### Single-nuclei RNA-seq analysis identifies distinct fibroblast and endothelial subpopulations mediating homeostasis, injury response, and mesenchymal-transition following distension

While sequencing-based ST approaches provided unique insights into the spatial context of genome-wide transcription, the technology remains limited in cellular resolution.^33^ Therefore, we further investigated the effects of distension at the single-cell level through single-nuclei RNA sequencing (snRNA-seq) analysis as a complementary approach. Using four paired cephalic vein samples from two canines, including two vein samples retrieved following distension with saline and two control vein samples (Table S1), we performed snRNA-seq analysis on a total of 1,188 nuclei derived from control veins and 1,223 nuclei derived from distended veins. Clustering of the nuclear transcriptomes yielded ten clusters visualized by UMAP embedding, representing endothelial, fibroblast, vascular smooth muscle, and myeloid cell types (Figure 4A) based on the expression of canonical markers (Figure S3). Owing to the plasticity of endothelial cells and fibroblasts, four distinct subpopulations also appeared within both the EC and FB clusters. Examining the distribution of these cell types and subpopulations showed an overall decrease in the proportion of ECs and SMCs following distension with an overall increase in FB and myeloid cell populations (Figure 4B). These alterations in cellular composition are noteworthy, as endothelial cell loss and increased immune cell infiltration, coupled with dedifferentiation and necrosis of contractile VSMCs, are associated with the early development of intimal hyperplasia in VGF.^20^

**Figure 4.**
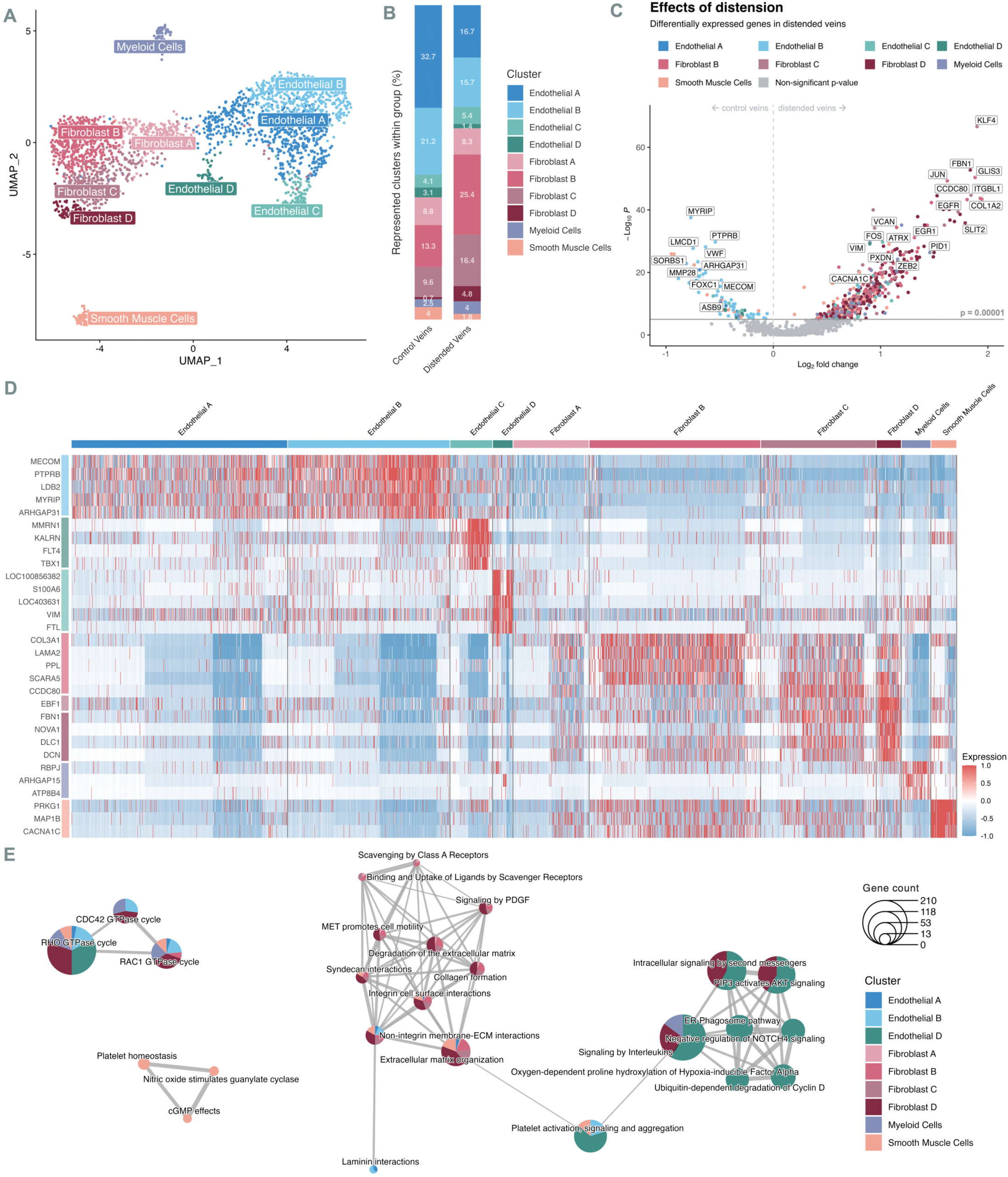
Single-nuclei transcriptomic analysis of control and distended veins. **(A)** UMAP displaying the clustering of nuclei derived from control and distended veins. Canonical cell types are based on the expression of marker genes (Figure S3). **(B)** Relative proportions of each cluster present within control vein and distended vein groups. **(C)** Volcano plot of differentially expressed genes within the distended vein compared to the control vein (*P* < 0.00001), wherein each gene is colored according to the cluster with the highest expression of the marker. **(D)** Heatmap of top marker genes associated with different cell types. Columns represent individual cells grouped by cell types, while rows display individual genes. Horizontal colored bars above the heatmap indicate the different cell types. Relative gene expression is shown in pseudo color, where blue represents low expression and red represents high expression. Top marker genes for clusters are defined by the average log_2_ fold-change of genes expressed in greater than 30% of nuclei within a cluster and *P* < 0.05. **(E)** Network of key differentially enriched pathways between clusters, derived from clusterProfiler analysis using the Reactome database (*P* < 0.05). Each node depicts the proportion of each cluster displaying enrichment of the associated pathway and the cumulative number of genes enriched in the pathway between the represented clusters.

Analysis of the effects of distension on the gene expression profile further reflected these differences in cellular composition (Figure 4B). Specifically, within distended veins, key transcription factors (TFs), including Krüppel-like factor 4 (*KLF4*), GLIS Family Zinc Finger 3 (*GLIS3*), early growth response factor 1 (*EGR1*), and zinc finger E-box-binding homeobox 2 (*ZEB2*), were all significantly upregulated (*P* < 0.00001), in addition to the AP-1 subunits, c-Fos (*FOS*) and c-Jun (*JUN*) (Figure 4C). While these TFs all have well-documented roles in cellular reprogramming and transdifferentiation, only *KLF4, EGR1, ZEB2,* and *AP-1* have been documented in vascular endothelial-to-mesenchymal and myofibroblast transitions. In contrast, *GLIS3* has not been previously reported to play a role in vascular remodeling, yet was found to be significantly upregulated in the Fibroblast B, C, and D clusters, as well as the Endothelial B and C clusters. Intriguingly, this TF was recently found to promote the transdifferentiation of fibroblasts into retinal pigmented epithelial cells,^34^ suggesting that it may play a significant role in fibroblast reprogramming observed in response to endothelial injury following distension.

In addition to these transcription factors, genes encoding ECM proteins, including the EGF-family proteins integrin subunit beta like 1 (*ITGBL1*), epidermal growth factor receptor (*EGFR*), fibrillin (*FBN1*), and versican (*VCAN*), as well as the secreted matrix proteins slit guidance ligand 2 (*SLIT2*), collagen I (*COL1A2*), peroxidasin (*PXDN*), and coiledcoil domain-containing protein 80 (*CCDC80*) were significantly upregulated (*P* < 0.00001) across the fibroblast subpopulations within the distended vein, highlighting the upregulation of ECM remodeling following distension (Figure 4D).

Notably, significantly downregulated genes (*P* < 0.00001) within the distended veins were primarily associated with maintaining endothelial identity and integrity, such as von Willebrand factor (*VWF*), vascular endothelial protein tyrosine phosphatase receptor type B (*PTPRB*),^35^ and MDS1 and EVI1 complex locus (*MECOM*)^36^ (Figure 4C). Regulators of endothelial injury response myosin VIIA and Rab interacting protein (*MYRIP*),^37^ forkhead box C1 protein (*FOXC1*),^38^ and LIM and cysteine-rich domains 1 (*LMCD1)*^39^ were also found to be significantly downregulated (*P* < 0.00001), particularly among the endothelial subpopulations (Figure 4C). Overall, the downregulation of these homeostatic and regulatory genes indicates that distension induces a transcriptomic shift towards the activation of ECs and the positive regulation of endothelial injury response.

Among the differentially expressed genes within the SMC population of distended veins, chromatin remodeler *ATRX* and VSMC-specific voltage-dependent calcium channel *CACNA1C* were significantly upregulated (*P* < 0.00001). Although the epigenetic role of ATRX has not been elucidated in the context of VSMC function, its upregulation among the SMC population may point toward the reprogramming of VSMCs following distension, as observed through the ST analysis. In contrast, the metalloprotease protein MMP-28 gene expression decreased in VSMCs following distension. Importantly, MMP-28 is considered a protective metalloprotease, wherein its depletion has been demonstrated to impede M2 macrophage activation, leading to impaired inflammatory response and cardiac dysfunction.^40^

We next explored the differentially expressed genes within each cluster (Figure 4D) and their associated pathways (Figure 4E), revealing similar trends between several cell types and subpopulations. First, Endothelial D, Fibroblast D, and myeloid cell clusters were significantly enriched (*P* < 0.05) for pathways related to cellular proliferation and migration, including AKT, NOTCH, and HIF signaling pathways, RHO and RAC GTPase cycle activation, and cell division, suggesting that these endothelial and fibroblast subpopulations represent activated phenotypes (Figure 4E). Upregulated genes among the fibroblast clusters overlapped in their enrichment of ECM remodeling pathways, integrin- and syndecan-mediated interactions, and MET-mediated cellular migration (*P* < 0.05), mirroring the pathways enriched within the media and adventitia observed via ST analysis (Figure 2B). Within the SMC cluster, gene expression was primarily associated with positive regulation of platelet homeostasis and negative regulation of proliferation and migration through nitric oxide modulation of cGMP, congruent with the increased proportion of this cluster observed in control veins compared to distended veins. However, similar to the contrasting SMC gene expression between control and distended veins observed through differential gene expression analysis (Figure 4C), several pathways, including ECM remodeling, RHO and RAC GTPase cycle activation, and platelet activation, were partially enriched (*P* < 0.005) within the SMC cluster. This data suggests that the portion of the SMC population associated with distended veins displays altered transcriptional regulation toward a synthetic phenotype following distension.

To elucidate the different subpopulations present among the fibroblast and endothelial cell clusters, we isolated both the fibroblast (Fibroblast A-D) and endothelial cell (Endothelial A-D) clusters, then conducted differential gene expression analysis between each cluster and identified key markers and biological themes differentiating each subpopulation (Figures 5–6). First, upon examining the effects of distension on the overall gene expression within the fibroblast population (Figure 5A), we found that downregulated genes in the fibroblast population after distension were primarily associated with the Fibroblast A subpopulation. These downregulated genes (*P* < 0.00001) overlapped with genes previously identified in control veins (Figure 4C) and also included the antiapoptotic regulator secreted frizzled-related protein 1 (*SFRP1*),^41^ the negative regulator of angiogenesis multimerin-2 (*MMRN2*),^42^ and negative regulator of vascular EC proliferation (*SELENOP*).^43^ In contrast, significantly upregulated genes (*P* < 0.00001) within the fibroblast population following distension included transcriptional promoters of cell proliferation and ECM proteins previously identified amongst the set of upregulated genes within the distended veins (Figure 4C), in addition to ECM-related genes laminin 2 subunit (*LAMA2*), dystonin (*DST*), and BicC family RNA binding protein 1 (*BICC1*). Interestingly, the roles of dystonin and BICC1 in vascular injury have not been previously investigated, but their function as an anchor within the actin cytoskeleton and negative regulator of Wnt signaling, respectively, suggest that they may play an important role in modulating fibroblast motility and transdifferentiation following vascular injury.

**Figure 5.**
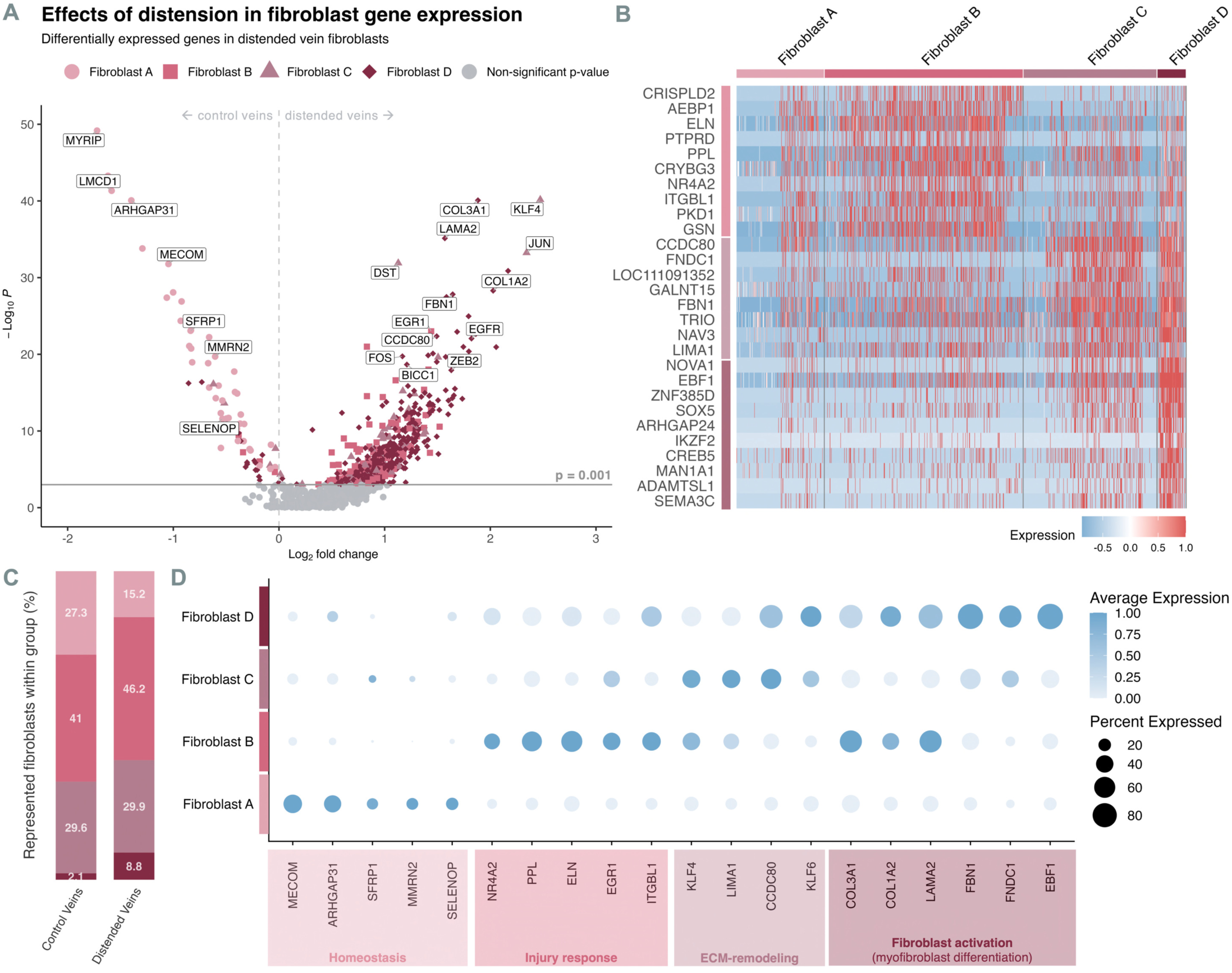
Characterization of fibroblast subpopulations. **(A)** Volcano plot of differentially expressed genes within fibroblast sub-clusters of the distended vein compared to the control vein (*P* < 0.001), wherein each gene is colored according to the fibroblast subpopulation with the highest expression of the marker. **(B)** Heatmap of top marker genes associated with different fibroblast sub-clusters. Columns represent individual cells grouped by cell types, while rows display individual genes. Horizontal colored bars above the heatmap indicate the different cell types. Relative gene expression is shown in pseudo color, where blue represents low expression and red represents high expression. Top markers for each fibroblast cluster are defined by the average log_2_ fold-change and *P* < 0.05. **(C)** Relative proportions of each fibroblast subpopulation present within control vein and distended vein groups. **(D)** Dot plot of the top markers representing the phenotype of each fibroblast subpopulation. The relative expression and percent of cells expressing specific markers are shown by shades of blue and the dot size, respectively.

**Figure 6.**
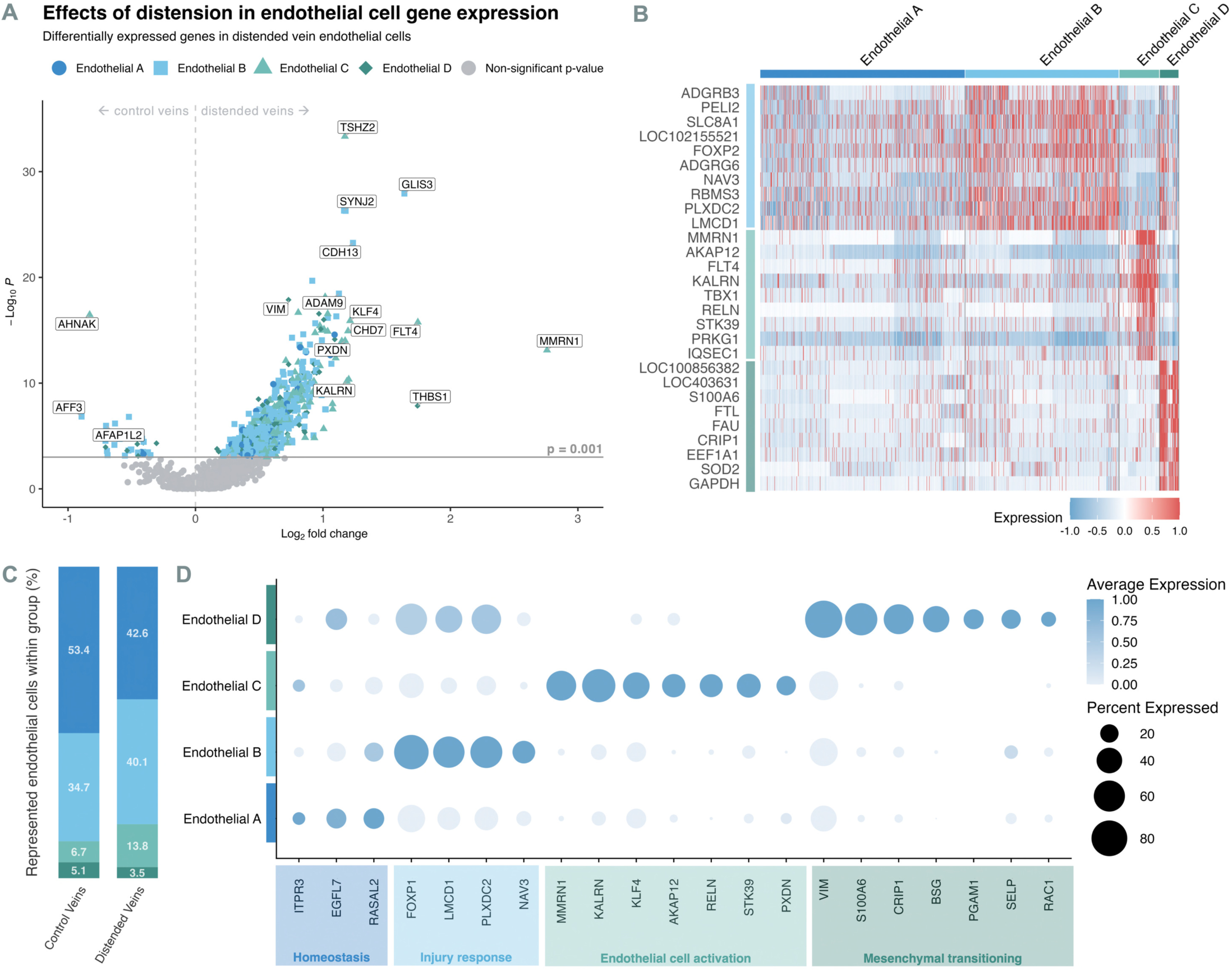
Characterization of endothelial cell subpopulations. **(A)** Volcano plot of differentially expressed genes within endothelial cells of the distended vein compared to the control vein, wherein each gene is colored according to the endothelial cell subpopulation with the highest expression of the marker. **(B)** Heatmap of top marker genes associated with different endothelial sub-clusters. Columns represent individual cells grouped by cell types, while rows display individual genes. Horizontal colored bars above the heatmap indicate the different cell types. Relative gene expression is shown in pseudo color, where blue represents low expression and red represents high expression. Top marker genes between endothelial clusters are defined by the average log_2_ fold-change and *P* < 0.05. **(C)** Relative proportions of each endothelial cell subpopulation present within control and distended vein groups. **(D)** Dot plot of the top markers representing the phenotype of each endothelial cell subpopulation. The relative expression and percent of cells expressing specific markers are shown by shades of blue and the dot size, respectively.

A closer look at the top gene markers associated with each fibroblast cluster (target fibroblast cluster versus other fibroblast clusters) facilitated the characterization of each fibroblast subpopulation (Figure 5B). First, expression of negative regulators of cell proliferation and migration (*ARHGAP31*, *SFRP1*, *MMRN2*, and *SELENOP*)^41–44^ within the Fibroblast A cluster and its increased proportion within control veins (Figure 5C) indicate that this fibroblast subpopulation is likely responsible for promoting vascular homeostasis following endothelial injury (Figure 5D). Similarly, within the Fibroblast B cluster (Figure 5B), the proapoptotic TF nuclear receptor 4A2 (*NR4A2*)^45^ was significantly upregulated with the profibrotic TF early growth response protein (*EGR1*), as well as the ECM proteins integrin subunit β-like 1 (*ITGBL1*), periplakin (*PPL*), and elastin (*ELN*) (*P* < 0.00001), signaling that these fibroblasts are likely involved in promoting fibroblast-mediated healing following endothelial injury (Figure 5D). In contrast, the Fibroblast C and D clusters both exhibited increased expression of genes associated with ECM remodeling; however, Fibroblast D specifically displayed significant upregulation of genes associated with myofibroblast transition (*P* < 0.00001), such as differentiation markers Krüppel-like factor 6 (*KLF6*) and EBF transcription factor 1 (*EBF1*), and production of type III collagen (*COL3A1*), type I collagen (*COL1A2*), laminin 2 subunit (*LAMA2*), fibrillin (*FBN1*), and fibronectin (*FN1*), indicating that Fibroblast C cells are responsible for ECM remodeling while Fibroblast D cells are protomyofibroblasts (Figure 5D). Collectively, the fibroblast subpopulations Fibroblast B (injury response fibroblasts), Fibroblast C (ECM remodeling fibroblasts), and Fibroblast D (protomyofibroblasts) were all enriched in their proportions within distended veins compared to control veins (Figure 5C), in addition to the overall increase in fibroblasts within the vein following distension (Figure 4B), revealing that distinct fibroblast subpopulations may play a crucial role in mediating the acute response to distension injury.

Within the endothelial cell subpopulations (Figure 6), many similarities appeared, as with the fibroblast population. However, distension notably resulted in the significant upregulation (*P* < 0.00001) of numerous genes within the endothelial population, including thrombospondin 1 (*THBS1*), multimerin 1 (*MMRN1*), and cadherins 7 (*CHD7*) and 13 (*CDH13*) (Figure 6A). Importantly, these genes have been reported to play significant roles in endothelial injury response and, moreover, the development of thrombosis, intimal hyperplasia, and atherosclerosis.^46–49^ The largest endothelial cluster, Endothelial A, was largely characterized by the downregulation of genes that were significantly overexpressed across the other endothelial clusters (Figure 6B). This cluster depicted an increase in proportion within control veins compared to distended veins (Figure 6C). Furthermore, this cluster displayed subtle expression of promoters of EC homeostasis, including inositol 1,4,5-trisphosphate receptor type 3 (*ITPR3*)^50^ and negative regulators of EC migration RAS protein activator like 2 (*RASAL2*) and EGF-like domain multiple 7 (*EGFL7*), suggesting that this subpopulation may be responsible for preventing EC activation and maintaining endothelial homeostasis (Figure 6D). Similarly, within the Endothelial B cluster (Figure 6B), genes previously implicated in mediating healing following endothelial injury were significantly upregulated (*P* < 0.00001), namely *LMCD1*, forkhead box protein P1 (*FOXP1*), plexin domain containing 2 (*PLXCD2*),^39,51,52^ and navigator-3 (*NAV3*). Intriguingly, navigator-3 was previously shown to suppress growth factor signaling, promote apoptosis, and decrease cell migration in epithelial cell lines and breast cancer models,^53^ and thus may play a similar role in maintaining endothelial homeostasis (Figure 6D).

In contrast to the Endothelial A and Endothelial B clusters, the Endothelial C and Endothelial D clusters exhibited markers of EC activation and EndMT, respectively. Specifically, the Endothelial C subpopulation (Figure 6B), which was the only endothelial population displaying overall enrichment in the distended vein (Figure 4B), exhibited significantly higher expression (*P* < 0.00001) of platelet adhesion ligand multimerin-1 (*MMRN1*)^47^ and mediator of activated EC migration A-kinase anchor protein 12 (*AKAP12*),^54^ as well as elevated expression of proliferation and migration genes: serine/threonine kinase 39 (*STK39*) and *KLF4*. Moreover, this subpopulation exhibited increased expression (*P* < 0.00001) of kalirin (*KALRN*), reelin (*RELN*), and peroxidasin (*PXDN)*; notably, kalirin promotes neointimal hyperplasia by activating RAC expression,^55^ while reelin increases leukocyte adhesion molecules ICAM-1, VCAM-1, and E-selectin and the progression of atherosclerosis,^56^ and peroxidasin has been correlated with vascular endothelial dysfunction by disrupting eNOS function.^57^ Together, this indicates that the Endothelial C subpopulation represents an activated EC phenotype expressing markers associated with poor prognosis following graft implantation (Figure 6D).

Lastly, the Endothelial D subpopulation displayed increased markers of EndMT (*P* < 0.00001), including vimentin (*VIM*), phosphoglycerate Mutase 1 (*PGAM1*), S100 calcium-binding protein A6 (*S100A6*), and cysteine-rich protein 1 (*CRIP1*) (Figure 6D). Intriguingly, while EndMT is often associated with vascular dysfunction, the proportion of this mesenchymal subpopulation was decreased in distended veins compared to control veins (Figure 6C), possibly reflecting the alternative role of mesenchymal cells as mediators of fibrosis and inflammation in proper vascular healing, such as that observed in type II epithelial-mesenchymal transition. Moreover, the concomitant increase in activated ECs (Endothelial C) and decrease in mesenchymal-transitioning cells (Endothelial D) in distended veins also supports recent findings that EndMT plays a marginal role in the contribution to neointimal formation leading to VGF and rather appears to represent an adaptive response capable of suppressing VSMC reprogramming and promoting quality vascular remodeling following graft implantation.^58^

### Integration of single-nuclei and spatial transcriptomics illustrates a significant impact of distension on the enrichment of activated endothelial and fibroblast subpopulations within the intima

To achieve the most granular view of the cellular transcriptome landscape, we integrated our single-nuclei and spatial transcriptomics data by deconvoluting the spatial voxels of control and distended veins based on the transcriptional signatures of the cellular subpopulations elucidated via snRNA-seq (Figure 7A).

**Figure 7.**
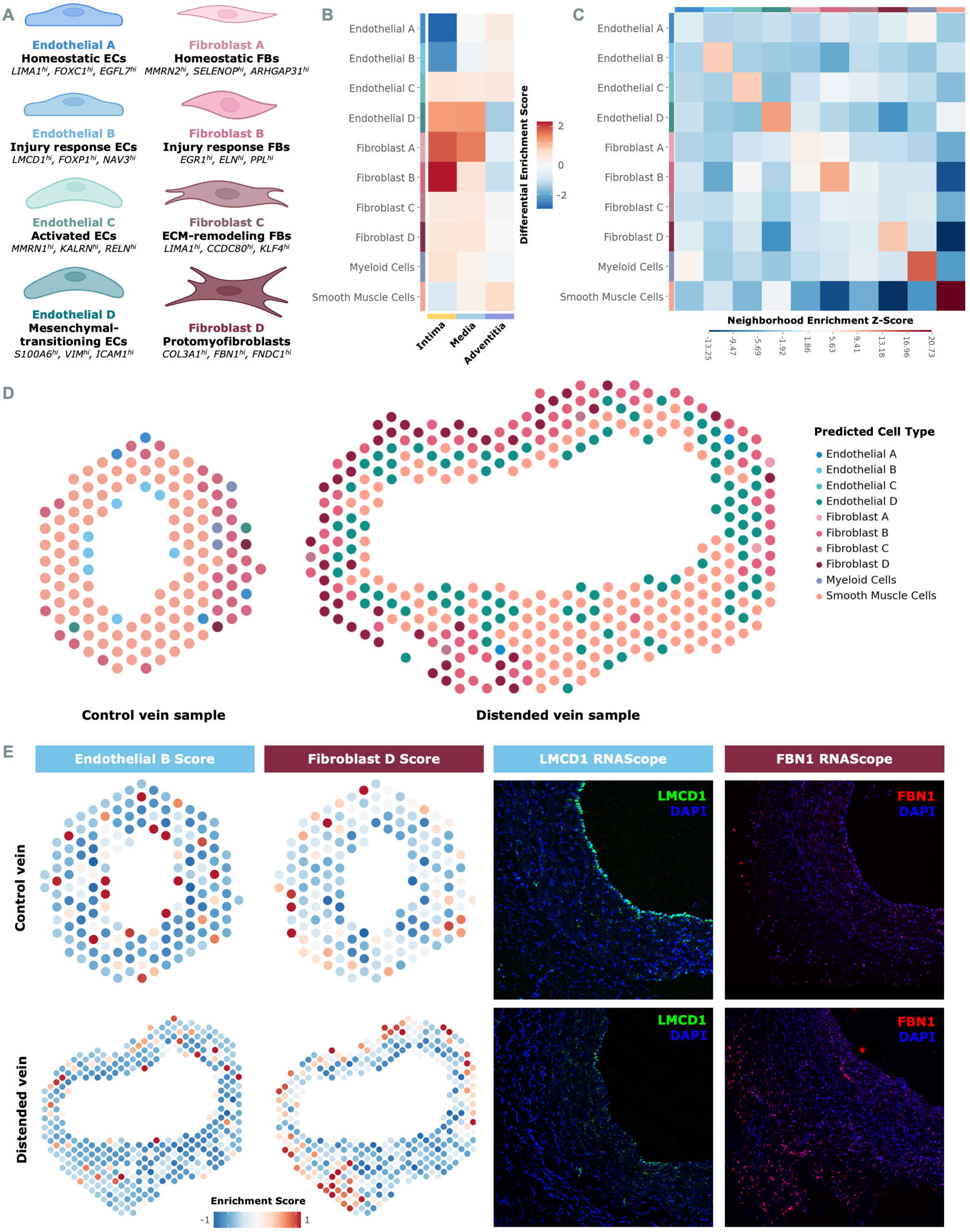
Deconvolution of spatial transcriptomic data through the integration of single-nuclei data. **(A)** Summary of distinct fibroblast and endothelial subpopulations identified in control and distended veins via snRNA-seq analysis along with expression ok key markers. **(B)** Heatmap displaying the differential enrichment scores of cellular subpopulations across layers of vein wall following distension. An increased differential score (red) indicates increased gene signature expression associated with the respective subpopulation in the distended veins relative to control veins, and blue indicates increased enrichment in control veins. **(C)** Heatmap of neighborhood enrichment Z-scores generated via Squidpy, illustrating the proximity of cellular populations to one another based on the dominant cell type assigned to each voxel. Red indicates enriched proximity between two populations, whereas blue indicates depleted proximity. **(D)** Spatial plots of representative control and distended vein samples displaying the dominant cell type within each voxel predicted by model-based deconvolution of the spatial transcriptomic data using the single-nuclei dataset via Cell2location. **(E)** Validation of the enrichment score (left) and model-based (panel D) integration approaches using RNAscope (right).

First, employing a scoring-based integration approach, we generated a set of representative and differentially expressed genes for each of the identified subpopulations, then utilized these gene signatures to assign an enrichment score of each cell type to the spatial voxels of control and distended veins (Method S3, Figure S4). To examine the spatial distribution of the subpopulations across the vein wall regions, we then averaged the cell type enrichment scores of voxels within the intima, media, and adventitia, generating enrichment score matrices for distended and control veins. Next, a differential enrichment matrix was computed to compare the relative spatial distribution of cell types between the distended and control veins, displaying corresponding increases and decreases in cell types across layers of the distended veins relative to control veins (Figure 7B). The resulting differential enrichment score matrix illustrated that Endothelial A and B subpopulations are reduced in the intimal layer of distended veins, pointing to decreased maintenance of EC homeostasis and mitigation of EC-mediated response at the site of the endothelial injury. Conversely, Endothelial C and D subpopulations appear enriched in the intima, reflecting the activation and dedifferentiation of ECs following distension. Furthermore, these mesenchymal-transitioning ECs (Endothelial D) were also enriched in the media of distended veins. Despite the overall decrease in the proportion of Endothelial D cells in distended veins captured by snRNA-seq (Figure 4B), the observed enrichment of the Endothelial D signature within the intima and media suggests that the limited Endothelial D subpopulation within distended veins may functionally differ from that of control veins, which were enriched in the adventitia. Increased fibroblast populations were also observed in the intima and media of distended veins, suggesting that fibroblasts may also localize to the site of endothelial injury following distension. Importantly, migration of fibroblasts to the intima plays a well-documented role in the development of intimal hyperplasia and atherosclerosis.^59^ Within the intima of distended veins, myeloid cells are also enriched, further underscoring the recruitment and infiltration of immune cells initiated by the effects of distension on the endothelium. Lastly, SMCs display increased enrichment in the adventitia of distended veins, supporting our hypothesis that while the overall proportion of SMCs decreases following distension, SMCs present in the distended veins may undergo VSMC-reprogramming toward a synthetic phenotype facilitating their migration to the adventitia.

As a complementary integration approach, we additionally performed model-based spatial deconvolution via Cell2Location with subsequent neighborhood enrichment analysis via Squidpy. Through neighborhood enrichment analysis, we examined the proximity probabilities of each cellular subpopulation in relation to one another throughout the vein (Figure 7C) based on the most abundant cellular subpopulation within each voxel (Figure 7D). This analysis showed the increased proximity of each subpopulation to other members of the respective subpopulation, particularly for SMCs, myeloid cells, and mesenchymal-transitioning cells (Endothelial D). This increased colocalization of the Endothelial D subpopulation aligns with the documented enrichment of intra-cell type signaling promoting EC activation and EndMT.^60^ Furthermore, the Endothelial D and SMC populations exhibited increased proximity, highlighting the crosstalk between ECs and VSMCs to mediate EC and VSMC activation in vascular dysfunction.^61^ To illustrate the spatial distribution of these cellular subpopulations, we constructed spatial plots displaying the most abundant population within each voxel, predicted using a Bayesian model generated from the snRNA-seq dataset via Cell2Location (Figure 7D). Congruent with the previous findings, the intima of the control vein displayed an increased Endothelial B (injury response) subpopulation, while the intima and media of the distended vein displayed enrichment of the Endothelial D population. Concurrent with this increase in Endothelial D abundance, a decrease in the overall proportion of VSMC-rich regions within the media is observed, mirroring the reduction in overall SMC proportion observed via single-nuclei analysis. Similarly, within the distended vein, enrichment of the protomyofibroblast (Fibroblast D) population is observed within the adventitia, demonstrating the increase in fibroblast activation following distension. To validate the integration of our single-nuclei and spatial datasets, we lastly performed RNAscope on control and distended vein samples (Figure 7E, Figure S5), using fluorescent antibodies for representative marker genes (*LMCD* and *FBN1*) corresponding to key subpopulations differentially enriched between control and distended veins (Endothelial B and Fibroblast D, respectively). The upregulation of *LMCD1* within the intima of control veins detected via RNAscope (Figure 7E, right) aligns with the predicted enrichment of the Endothelial B population within the intima of control veins observed by our scoring-based (Figure 7E, left) and model-based (Figure 7D) integration approaches. Similarly, the upregulation of *FBN1* within the adventitia of distended veins (Figure 7E, right) aligns with the predicted enrichment of the Fibroblast D population within the adventitia of distended veins, also observed by our integration approaches.

Taken together, the results of the integrated single-nuclei and spatial transcriptomic analysis elucidate the effects of distension on the cellular architecture of the vein. Most notably, the enrichment of activated ECs within the intima and migratory EndMT cells extending toward the media is observed following distension, along with the migration of fibroblasts to the intima and a decrease in homeostatic and injury response in luminal ECs. Importantly, these changes are also implicated in the development of IH and atherosclerosis, further suggesting that distension may initiate the pathways implicated in these VGF-related pathologies.

### Cellular communication analysis elucidates key signaling molecules promoting inflammation and vascular injury responses following distension

Finally, to elucidate the intercellular communication network between the subpopulations present in the veins following distension, we analyzed the number and strength of incoming and outgoing signaling interactions for each cell type, as well as the key signaling molecules mediating these interactions, via cellular communication analysis.^62^ Overall, the total number of intercellular signaling interactions within distended veins increased in addition to the strength of these interactions (Figure S6). Most notably, signaling to the Endothelial B (injury response ECs), Endothelial D (mesenchymal-transitioning ECs), Fibroblast C (ECM remodeling FBs), Fibroblast D (protomyofibroblasts), myeloid cells, and SMCs increased significantly following distension (Figure 8A-B). Furthermore, the strength of signaling to the mesenchymal-transitioning and myeloid cells in distended veins greatly increased (Figure S6), highlighting the roles of these populations in response to distension.

**Figure 8.**
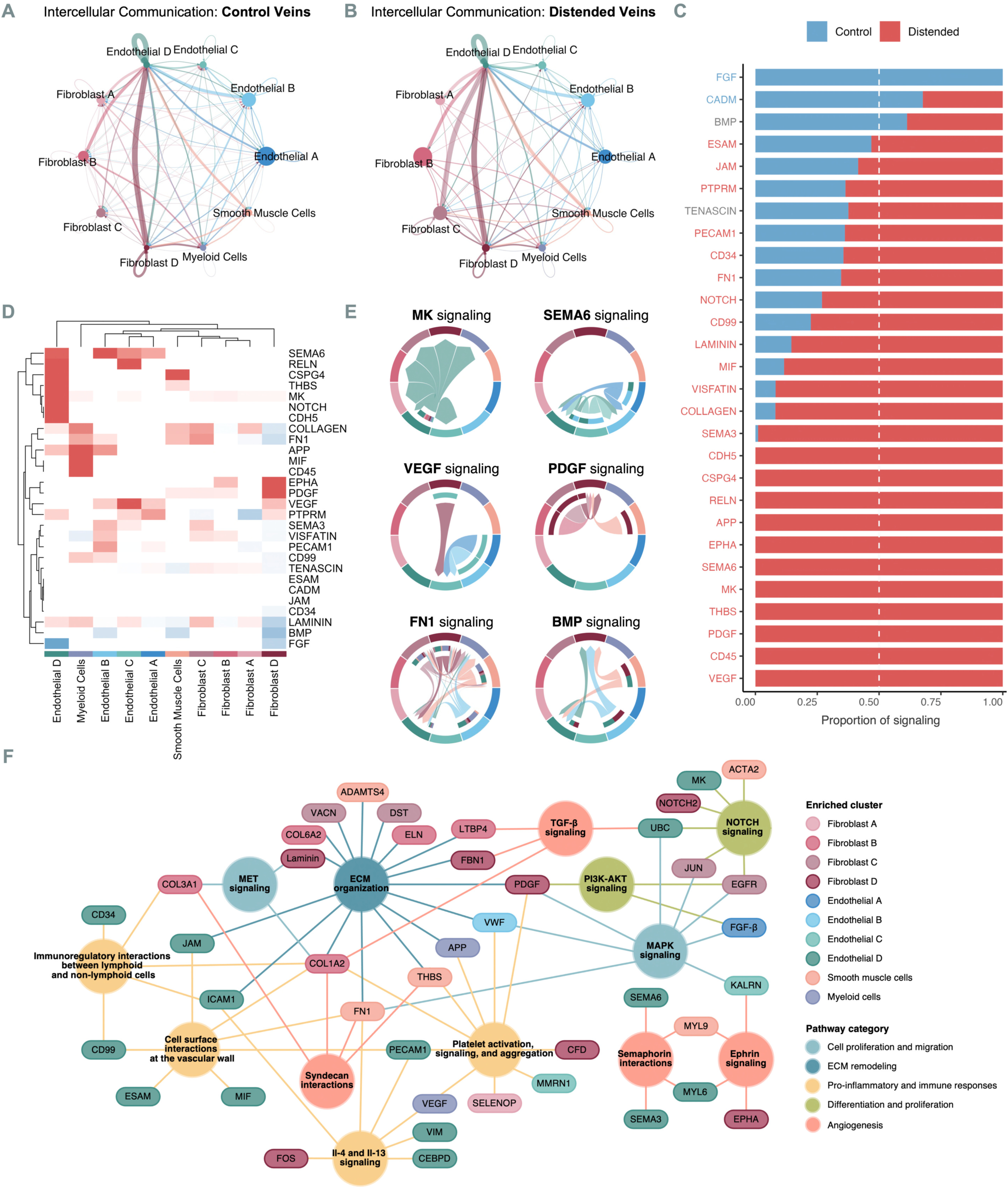
Intercellular communication analysis. Network displaying the number of intercellular communication interactions between subpopulations for control **(A)** and distended veins **(B)**. **(C)** The proportion of key signaling molecules active in control versus distended veins. **(D)** Heatmap of the differential signaling pattern (both incoming and outgoing signaling per cell type or subpopulation) in distended veins (red) compared to control veins (blue). **(E)** Chord diagrams illustrate intercellular communication networks for distended veins, with the exception of BMP, which is illustrated for the control vein. The outer ring and arrows of the diagram represent the signaling produced by each cellular subpopulation, while the inner ring represents which subpopulations are receiving the outgoing signals. **(F)** Network depicting the interactions of key signaling molecules and differentially expressed genes across pathways implicated in distension injury.

To characterize the intercellular communication among these subpopulations, we first examined which signaling pathways were enhanced in distended versus control veins (Figure 8C), revealing several pathways also observed in our ST and single-nuclei analyses, including ephrin (EPHA), reelin (RLN), collagen, laminin, and fibronectin (FN1). In addition to these previously implicated pathways, several interesting signaling pathways emerged. First, within the control veins, signaling pathways associated with vascular healing and angiogenesis were enriched, namely fibroblast growth factor (FGF),^63^ cell adhesion molecule (CADM),^64^ and bone morphogenic protein (BMP)^65^ mediated signaling pathways. In contrast, distended veins were enriched for disparate angiogenic signaling pathways, particularly those mediated by vascular endothelial growth factors (VEGF), platelet-derived growth factors (PDGF), midkine neurite growth-promoting factor 2 (MK), and semaphorins (SEMA6, SEMA3). Notably, the upregulation of VEGF, PDGF, and MK have all been well-documented as key contributors to intimal hyperplasia, contributing to graft failure.^66–68^

Next, to illuminate which subpopulations are modulating these key signaling pathways in distended veins compared to control veins, we generated a heatmap displaying the differential signaling of each pathway within the cellular subpopulations (Figure 8D) and elucidated the intercellular communication networks mediated by these pathways (Figure 8E). Markedly, signaling within the Endothelial D (mesenchymal-transitioning ECs) and Fibroblast D (protomyofibroblasts) subpopulations were most significantly affected by distension, closely followed by the Endothelial C (activated ECs) and SMC populations.

Within the Endothelial D (mesenchymal-transitioning ECs) population, signaling pathways associated with cell growth and differentiation (MK, NOTCH, CSPG4),^69,70^ neointimal formation and hyperplasia (THBS),^46^ and leukocyte adhesion and atherosclerosis (RELN)^56^ were specifically enhanced following distension. Similarly, within the SMC population, signaling associated with VSMC activation, namely chondroitin sulfate proteoglycan 4 (*CSPG4*) and fibronectin (*FN1*) pathways, were enriched, indicating that VSMC-reprogramming is initiated by distension injury. Inspecting the intercellular communication networks for these pathways unveiled that Endothelial D and SMC populations are primarily responsible for producing ligands associated with these pathways, whose surface receptors were displayed on other EC and FB subpopulations. For example, the outgoing MK signal (*MDK*) produced by the Endothelial D population interacted with the low-density lipoprotein (*LPR1*), nucleolin (*NCL*), and integrin (*ITGA6*, *ITGB1*) receptors present across the four fibroblast populations, activated EC subpopulation (Endothelial C), and myeloid cells, suggesting a potential role of the Endothelial D subpopulation in promoting the proliferation of fibroblasts, activated ECs, and immune cells following distension (Figure 8E). Likewise, the SMC population of the distended veins predominantly produced fibronectin (*FN1*), which was received by integrin surface receptors (*ITGA5*, *ITGB1*) on Endothelial D cells, as well as CD44 receptors on Fibroblast C (ECM remodeling FBs), Fibroblast D (protomyofibroblast), and myeloid cells (Figure 8E); stimulation by fibronectin is known to promote EndMT and myofibroblast transdifferentiation, suggesting that differentiation of the cell types is strongly influenced by the SMC population within distended veins. Collectively, these findings further support our hypothesis that while the proportions of mesenchymal-transitioning ECs and VSMC populations are decreased in distended veins, the residing populations of these cell types are phenotypically different from those of the control veins, promoting cell proliferation, migration, and activation that may initiate the development of intimal hyperplasia in VGF.

Within the Fibroblast D (protomyofibroblasts) population, increased signaling via the ephrin (EPHA), platelet-derived growth factor (PDGF), and vascular endothelial growth factor (VEGF) pathways were observed (Figure 8D). These pathways all have well-documented roles in angiogenesis, including the implication of VEGF and PDGF in the development of intimal hyperplasia.^66,67^ Interestingly, ephrin signaling in the distended vein was mediated by ephrin-A5 (*EFNA5*), produced by Fibroblast D cells, with ephrin-A3 (*EFNA3*) receptors displayed on both Fibroblast D and Fibroblast B (injury response FBs) cells. While ephrin-B has been thoroughly investigated for its role in angiogenesis as a VEGF receptor,^71^ the role of ephrin-A signaling remains understudied and may warrant future investigation. Notably, VEGF signaling was also upregulated among the Endothelial C (activated ECs) population, which was uniquely enriched in distended veins. A closer examination of the VEGF signaling network subsequently revealed that VEGF signaling is mediated by VEGFC, produced by Fibroblast D, Endothelial A (homeostatic ECs), and Endothelial B (injury response ECs) cells, and received exclusively by VEGFR3 receptors on the Endothelial C subpopulation (Figure 8E). This finding suggests that the Fibroblast D, Endothelial A, and Endothelial B cells induce the activation of ECs in response to distension. Likewise, PDGF signaling via the PDGFD ligand was produced by Fibroblast D, Fibroblast C (ECM remodeling FBs), Fibroblast B (injury response FBs), and VSMC populations and received exclusively by Fibroblast D cells by PDGFRB expression, facilitating the activation of these fibroblasts following distension.

Together, the results of our intercellular communication analysis provide greater insight into the distinct roles of specific EC and FB subpopulations in combination with VSMCs and immune cells in modulating the response to endothelial injury initiated by distension. Combining the key signaling molecules with prominent differentially expressed genes within distended veins, we generated a network of the key pathways implicated in distension injury, illustrating the effects of distension on genomic expression within the vein (Figure 8F). The results of these analyses cumulatively demonstrate that distension elicits a coordinated response throughout the vein wall, driving the activation of endothelial cells, fibroblasts, and vascular smooth muscle cells and stimulating proinflammatory and immune responses.

### Vascular remodeling, proinflammatory pathways, and key genes (FBN1, VCAN, GLIS3) initiated by distension persist post-bypass

Lastly, to determine the relation of the above findings to the pathophysiology of vein grafts post-implantation, we extended our investigation to a vein graft sample retrieved 24 hours post-bypass, examining the cellular composition and transcriptomic profile. At 24 hours post-bypass, an increase in myeloid cell, protomyofibroblast, injury-response EC, and mesenchymal-transitioning EC subpopulations was observed with a concomitant decrease in homeostatic ECs and fibroblasts (Figure 9A). These findings are congruent with a response to increased cellular activation, proinflammatory, and immune signaling pathways upregulated at the time of distension. Furthermore, spatial transcriptomic analysis of the implanted graft illustrated the distribution of these subpopulations with protomyofibroblasts extending from the adventitia to the media and injury response and EndMT ECs throughout the intima (Figure 9B). Next, to examine how the genomic response initiated by distension extends to the transcriptomic profile observed 24 hours post-implantation, we constructed a temporally-resolved multilayer gene network that illustrates the trajectory of the genomic response from control veins to distended and implanted veins. Notably, there were several significantly upregulated genes at the time of distension, including fibrillin-1 (*FBN1*) and versican (*VCAN*),^72,73^ that were subsequently driving the expression of genes implicated in vascular remodeling and graft failure, such as *IL-6*, *TGFBR1*, *SMAD4*, and *ADAMTS9*,^74–76^ within the implanted vein (Figure 9C). Additionally, novel genes of interest identified through the above single-nuclei transcriptomic analysis, such as *GLIS3*, were present among the drivers of post-implantation gene expression, underscoring the advantage of this approach in identifying potential therapeutic targets. Specifically, the transcription factor *GLIS3*, which was significantly upregulated at the time of distension and has been previously demonstrated to play a role in epithelial cell reprogramming,^34^ was found to drive expression of TAZ (*WWTR1*), a central regulator in HIPPO signaling pathway driving EndMT,^77^ within the grafted vein (Figure 9C). Finally, to validate the expression of these genes at the protein level, we performed IHC analysis on control, distended, and grafted vein samples, staining for the expression of VCAN, FBN-1, and GLIS3 (Figure 9D, Figure S7). Mirroring the results of transcriptomic analyses, an increase in the expression of these proteins was observed at the time of distension, which was further increased 24 hours post-implantation by IHC staining.

**Figure 9.**
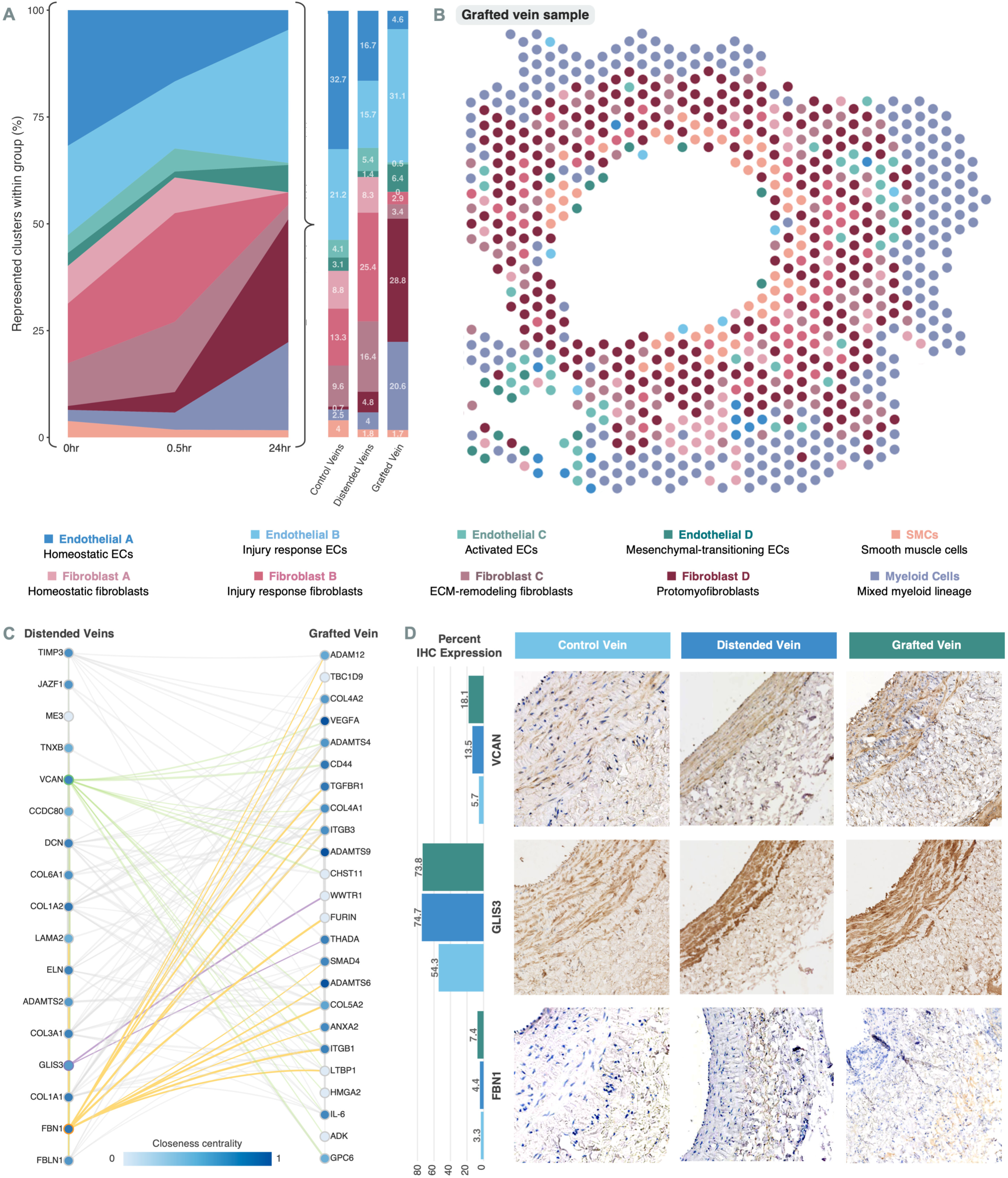
Relationship of pre- and post-implantation transcriptomic profiles and select protein expression. **(A)** Relative proportions of each fibroblast subpopulation present within control vein and distended vein groups. **(B)** Spatial plot of grafted vein sample displaying the dominant cell type within each voxel based on deconvolution of the spatial transcriptomic data using the single-nuclei dataset via Cell2location. **(C)** Temporally resolved multilayer gene network analysis illustrating the effects of key genes (circled) at the time of vein harvesting and distension (left) to 24 hours post-implantation (right). **(D)** Immunohistochemistry staining of control, distended, and grafted veins with VCAN, GLIS3, or FBN with quantification values (left).

## Discussion

Vein bypass graft failure and thrombosis following peripheral and coronary artery bypass grafting can result in limb loss, heart failure, and death. Thus, to overcome this challenge, there is a need to elucidate the molecular mechanisms underlying graft failure and identify potential targets for therapeutic development. VGF is primarily attributed to IH, a process that causes thickening and intraluminal narrowing of a bypass graft, obstructing blood flow and ultimately leading to graft thrombosis.^78^ It has been posited that IH may occur in response to bypass implantation, where repeated exposure to arterial hemodynamics, ischemia-reperfusion injury, and biomechanical stressors results in damage to the endothelium, triggering changes in vascular gene expression and concomitant transition of quiescent ECs to an activated proinflammatory and prothrombotic state and VSMCs from contractile to an activated synthetic state.^78–81^ Subsequent VSMC proliferation and migration, coupled with excessive ECM production and fibroblast migration, ultimately generates obstructive IH lesions and VG atherosclerosis, contributing to graft failure. While the development of IH following graft implantation has been investigated, we have sought to elucidate the onset of these maladaptive pathways, beginning at the time of graft harvesting and its connections with the changes seen post-implantation. Through this study, we unveiled that conduit distension during graft harvesting induces endothelial injury and initiates the dysregulation of genes and pathways implicated in graft failure, which partially drives injury-related pathways identified following implantation. Harnessing an integrated single-nuclei and spatial transcriptomics approach, we provide unprecedented insights into the genomic effects of distension, the unique cellular subpopulations mediating this response, and the spatial heterogeneity of acute distension injury.

By leveraging recent advances in spatial transcriptomic technologies, we performed the first reported spatial transcriptomic investigation of vein tissue samples. This methodology uniquely facilitates unbiased genomic analysis with spatial resolution, capturing the expression of over 35,000 genes localized across the layers of the vein wall. Through this approach, we illustrated the effects of distension on the genomic and cellular landscape of the vein graft. Aligning with previous investigations of IH in VGF,^78^ we found that distension alters gene expression heterogeneously throughout the layer of the vein, reflecting the traumatic sheer stress resulting from distention to diameters greater than the existing cellular architecture permits. Specifically, examining each layer individually, we identified markers of EC, VSMC, and fibroblast activation within the intima, media, and adventitia, respectively. Among these markers, we identified the upregulation of several genes implicated in IH and atherosclerosis, including fibronection (*FN1*) within the intima and vimentin (*VIM*) within the adventitia.^82,83^ Concomitant with the differential expression of these genes in distended veins, we identified corresponding enriched pathways within the intima, media, and adventitia; pathways associated with ECM remodeling, cell proliferation, and migration (collagen, integrin, syndecan, and MET signaling) were upregulated in the intima and media, while platelet activation pathways were significantly upregulated in the media and profibrotic IL-4 and IL-13 signaling were upregulated in both the adventitia and media. To validate these findings, we further performed unsupervised clustering of the spatial voxels based on their transcriptomic profiles, identifying clusters that heterogeneously localized to distinct regions of the vein. Interestingly, we identified a cluster (I1) exhibiting significant upregulation of markers of EC activation and EndMT, including fibronectin (*FN1*), VWF, tropomyosin 1 (*TPM1*), α-SMA (*ACTA2*), γ-SMA (*ACTG1*), and thrombospondin 1 (*THBS1*), which protruded from the intima into the media of the distended vein. Furthermore, within distended veins, we identified enriched medial and adventitial clusters displaying markers of cell activation and migration, including MAPK and L1CAM signaling within the media, and markers of inflammation, including IL-4 and IL-13 signaling in the adventitia. Collectively, our spatial transcriptomic analyses reveal that distension results in the upregulation of ECM remodeling, cellular activation, and inflammatory pathways across the intima, media, and adventitia of the vein, highlighting the sensitivity of veins to manipulation and dissection. While these pathways play critical roles in the physiological response to endothelial injury, their dysregulation leads to IH development. Therefore, the upregulation of these at the time of distension may illustrate the initiation of pathways that later become dysregulated and lead to IH in the context of graft failure. Vasodilatory agents have been suggested as potential therapeutics to reduce endothelial injury, but even with the use of papaverine-based solution for intraluminal vasodilation in this study, distension was found to impart significant endothelial injury, underscoring the need for alternative approaches.

These findings are further supported by the results of our single-nuclei transcriptomic analysis, wherein we identified distinct populations of ECs, FBs, VSMCs, and immune cells mediating the genomic response to distension. Overall, we observed decreased EC and VSMC populations following distension, with a concurrent increase in FB and myeloid cell populations. We posit that these changes to the cellular architecture reflect injury and desquamation of the endothelial layer. Moreover, EC loss, leukocyte recruitment, and dedifferentiation of contractile VSMCs are documented in the early phases of IH development in VGF,^20^ supporting the idea that IH leading to VGF may be initiated during graft harvesting. Subsequent analysis of the nuclear transcriptomes further enabled the identification and characterization of unique endothelial cell and fibroblast subpopulations associated with maintaining vascular homeostasis (Endothelial A, Fibroblast A), endothelial injury response and repair (Endothelial B, Fibroblast B), ECM remodeling (Fibroblast C), and cellular activation (Endothelial C) and transdifferentiation (Endothelial D, Fibroblast D). Following distension, the proportion of injury response and activated ECs increased among the EC population, and within the FB population, the proportion of injury response FBs, ECM remodeling FBs, and protomyofibroblasts increased. Intriguingly, the proportion of mesenchymal transitioning ECs (Endothelial D) was increased in the control veins compared to distended veins despite the documented role of EndMT in vascular dysfunction. Notably, this finding supports the recent work by Wu and coworkers, which illuminated the pro-fibrotic and pro-inflammatory role of mesenchymal cells in proper wound healing,^58^ similar to that of type II epithelial-mesenchymal transition. Furthermore, differential gene expression, pathway enrichment, and intercellular communication analyses together suggested that the EndMT (Endothelial D) subpopulation exhibits a disparate transcriptomic profile in distended veins compared to control veins. Specifically, Endothelial D cells within the distended veins were upregulated in cell growth, proliferation, and migration (MK, CSPG4), leukocyte adhesion and stenosis (RELN), and neointimal formation and hyperplasia (THBS) associated pathways. While the population of VSMCs was reduced in distended veins, the residing VSMCs of distended veins exhibited a distinct expression profile compared to those of control veins, wherein the VSMCs of distended veins produced greater collagen and fibronectin, promoting EndMT and transdifferentiation-associated pathways. Additionally, through the single-nuclei transcriptomic analyses, we also identified the upregulation of several key genes previously implicated in VGF, including versican (*VCAN*), fibrillin (*FBN1*), and vascular endothelial growth factor-C (*VEGFC*). Moreover, we unveiled several novel genes of interest that were significantly upregulated following distension and have yet to be investigated in the context of graft implantation and failure, including neuron navigator 3 (*NAV3*), semaphorin-6A (*SEMA6A*), BicC family RNA binding protein 1 (*BICC1*), and dystonin (*DST*). Furthermore, owing to the increased resolution of nuclear transcripts via single-nuclei analysis, we also identified significant upregulation of several key transcriptional regulators following distension, including Krüppel-like factor 4 (*KLF4*), early growth response factor 1 (*EGR1*), and zinc finger E-box-binding homeobox 2 (*ZEB2*), and GLIS Family Zinc Finger 3 (*GLIS3*), the latter of which has previously been implicated in fibroblast to epithelial cell transdifferentiation^34^ but warrants investigation for its role in regulating transdifferentiation during vascular remodeling. Taken together, the results of our single-nuclei analysis elucidate differentially expressed key genes and enriched pathways in distended veins, underscoring the acute effects of distension and providing a foundation for future analyses of these mechanisms. Integrating the ST and snRNA-seq datasets, we illustrated that EndMT-associated (Endothelial D) gene expression was increased in the intima and media of distended veins, along with homeostatic and injury response FB subpopulations and myeloid cells. Collectively, these findings illustrate that distension results in acute injury to the endothelium, triggering EC, VSMC, and FB activation with concomitant increases in the associated pathways involving cell proliferation and migration, ECM remodeling, inflammatory and immune responses, and transdifferentiation. This switch from quiescent to activated, pro-inflammatory phenotypes, coupled with the upregulation of prothrombogenic factors such as PDGF, Tumor Necrosis Factor (TNF), cytokines, and other damage-associated molecular patterns (DAMPs), as part of the initial responses to injury contributing to VGF, has long been noted.^84,85^ However, through this work, we have unveiled that these mechanisms may originate during the initial vein harvest dissection and distention process, highlighting the potential advantage of alternative harvesting techniques or early gene-targeted therapeutic interventions in improving vein graft patency.

Significantly, the pro-inflammatory changes identified immediately following distension, both in terms of cellular population and gene expression, persist post-implantation and, in some instances, appear to be driving graft failure mechanisms in the implanted vein. One of the key mechanisms of pathological intimal hyperplasia formation is excessive myofibroblast differentiation from quiescent fibroblast and migration from adventitia to the neointima.^86,87^ This excessive myofibroblast migration to the neointima has been previously implicated in graft failure following implantation,^20,88,89^ and the increase in protomyofibroblast seen in our samples 24 hours post-implantation indicates a similarly prominent role in the cellular response to injury post-implantation. However, while this phenotypical shift in fibroblasts has been primarily attributed to the arterialization of the vein following implantation,^78,90^ our results indicate that this transition is initiated during harvest and distension of the vein, as demonstrated by the increase in the proportion of injury response and ECM-remodeling FBs and protomyofibroblasts following vein distension, as well as their migration to the intima illustrated by spatial transcriptomics. Additionally, our temporally-resolved multilayer gene network showing the relationship between the distended and post-implantation transcriptomic profiles indicates that several genes previously implicated in both early and late graft failure are present at 24 hours post-implantation and driven by the upregulation of key genes during graft distension. Namely, the upregulation of IL-6, responsible for promoting leukocyte adhesion and vascular wall infiltration leading to thrombosis, and TGF-ß receptor 1, which plays a key role in mediating SMC differentiation as part of the TGF-ß signaling pathway, were demonstrated to be directly influenced by upregulated genes at the time of distension, including fibrillin-1 (*FBN1*) and versican (*VCAN*),^91–93^ which were validated by IHC. These findings further illustrate the putative role of vein harvest and distension as the initial insult that results in the cascade of physiopathological processes leading to VGF and highlights pre-implantation as a promising time point for intervention.

While the study presented herein pioneers the investigation of venous pathophysiologies, specifically acute distension injury through an integrated spatial and single-nuclei transcriptomics approach, it has several limitations. First, the study was not conducted in human vessels, and although we have previously utilized canine end-to-side cephalic vein interposition VG bypass models to investigate VGF after bypass grafting due to the animal model’s established relevance to pre-clinical human disease of IH and VG atherosclerosis, the difference in injury pathways between canine and human may exist.^94^ Additionally, as this dataset does not include failed graft samples, we cannot determine whether the genomic changes demonstrated in our work would ultimately result in graft failure. Furthermore, the study sample size limits the statistical power of our results, specifically when comparing cellular populations between distended and control veins. Population comparisons are also inherently confounded by the ability of different transcriptomic assays to capture specific cell types. For example, through our investigation, we found that the ST methodology was superior in capturing SMC-associated gene expression, while snRNA-seq provided high resolution of EC and FB population gene expression. While this limitation can impair statistical significance, it also underscores the utility of an integrated spatial and single-nuclei transcriptomics approach for efficiently capturing the heterogeneous architecture of the vein.

Conduit distension during vein graft harvesting is canonically performed during peripheral artery bypass surgery to prevent vasospasms and ensure adequate vein graft diameter and compliance.^95^ During this process, the saphenous vein is exposed along its entire length, followed by the insertion of a cannula at the distal aspect of the vein and subsequent distension. However, the sheer stress imparted on the vascular wall from elevated distension pressure has been illustrated to cause damage to the endothelium and media.^95^ Thus, to circumvent these effects, recent efforts to minimize manipulation and avoid distention of the vein graft have been employed, including the use of warmed papaverine solution to vasodilate the vein graft during distention. This process has been shown to reduce endothelial loss by scanning electron microscopy.^96,97^ Furthermore, in-situ vein bypass techniques, wherein the vein graft is left along its anatomic course without complete explanation, were performed to minimize dissection and manipulation of the vein graft. However, this type of in-situ bypass can only be utilized for certain distal arterial targets and has not been found to be superior to previously described techniques.^98^ Others have advocated for a “no-touch” vein harvest technique, in which the adventitia and perivascular tissue of the vein are kept intact during dissection while employing minimal to no distension.^99^ This “no-touch” harvest technique has been recently popularized in the cardiac literature for vein harvest during coronary artery bypass grafting, with a randomized control trial showing a significantly reduced risk of VG occlusion at one year.^8^ The results of our study support the use of these alternative graft harvesting approaches, highlighting the adverse effects of distensions on the vein and providing a possible explanatory mechanism for the significant improvement in graft patency seen in grafts implanted with minimal distension compared to those harvested with the conventional technique.

In conclusion, the results of this work present the first investigations of the acute genomic responses to distension during vein graft harvesting by leveraging single-nuclei and spatial transcriptomics. These responses, mediated by distinct populations of endothelial, fibroblast, smooth muscle, and myeloid cells throughout the layers of the vein, initiate EC, FB, and SMC activation through upregulation of cell proliferation, differentiation, and migration, ECM remodeling, proinflammatory, and angiogenic pathways. Furthermore, the upregulation of genes implicated in IH, endothelial dysfunction, and atherosclerosis, all contributing to VGF, were observed following graft distension, in addition to novel genes of interest that may play a role in VGF. As a whole, this work provides a foundation for future investigations of vein graft harvesting, graft implantation, and graft failure, exemplifies the utility of spatial and single-nuclei transcriptomics for the investigation of vascular pathologies, and identifies potential targets for the development of therapeutics that limit graft failure.

**Novelty and Significance**

**What Is Known?**

▪ Autogenous vein, namely the greater saphenous vein, is the most widely used conduit for coronary artery bypass grafting (CABG) and is the gold-standard conduit for peripheral arterial bypass surgery, however, the 1-year primary patency rate of coronary and peripheral vein grafts is as low as 60%, primarily due to maladaptive vascular wall remodeling response and the development of intimal hyperplasia (IH).
▪ While the late mechanisms leading to vein graft failure (VGF) have received great attention, less attention has been given to how these mechanisms may be initiated, particularly the role of graft harvesting techniques in the upregulation of maladaptive pathways implicated in VGF.

**What New Information Does This Article Contribute?**

▪ This study elucidated the acute genomic response following distension during vein graft harvesting, including the upregulation of genes and pathways previously implicated in VGF following implantation, and lays a foundation for future single-nuclei and spatial transcriptomics approaches to studying the mechanisms of vein graft failure and other vascular pathologies.
▪ Leveraging single-nuclei and spatial transcriptomics, this work unveils the mechanisms of endothelial cell (EC), fibroblast (FB), and vascular smooth mucle cells (VSMC) activation following distension via the upregulation of cell proliferation, differentiation, and migration, ECM remodeling, proinflammatory, and angiogenic pathways by distinct populations of ECs, FBs, VSMCs, and myeloid cells.

This work sheds light on the acute genomic effects of distension during vein graft harvesting, in which EC, FB, and VSMC activation is initiated through the upregulation of cell proliferation, differentiation, and migration, ECM remodeling, proinflammatory, and angiogenic pathways. We identified distinct subpopulations of Ecs, FBs, and VSMCs mediating this response, which were localized to specific regions of the vascular wall following distension, highlighting the effects of the traumatic sheer stress via distension on the pathophysiology of the vein. Furthermore, our analyses collectively suggest that the mechanisms implicated in VGF, including IH and endothelial dysfunction that are seen post-implantation, may originate at the time of distension during graft harvesting, underscoring the underscoring the importance of careful vein harvesting techniques and providing a basis for designing gene-targeted therapies to minimize maladaptive distention injury responses.

## Article Information

## Acknowledgments

The RNA sequencing in this work was supported by the Emory Integrated Genomics Core at Emory University, subsidized by the School of Medicine and Winship Cancer Institute. Dr. Vasantha L. Kolachala (Department of Pediatrics, Emory University) aided in the Visium assay standardization. Vaunita Parihar performed H&E imaging of the Visium slides and immunohistochemistry staining at the Cancer Tissue and Pathology shared resource of Winship Cancer Institute of Emory University and NIH/NCI under award number P30CA138292. The content is solely the authors’ responsibility and does not necessarily represent the official views of the National Institutes of Health. Figure illustrations were made with BioRender.com.

## Supplemental Materials

Methods S1-S5

Tables S1

Figures S1-S7

