## Supplemental Materials for "Integrated single-nuclei and spatial transcriptomic analysis reveals propagation of early acute vein harvest and distension injury signaling pathways following arterial implantation"

Manoj K. Bhasin, MS, PhD

Aflac Cancer and Blood Disorders Center

Children Healthcare of Atlanta

Woodruff Memorial Research Building, Room 4107

101 Woodruff Circle, 4th Floor East

Emory School of Medicine

Atlanta, GA 30322

### Table of Contents

#### I. Supplemental Methods

**Method S1.** Annotation of spatial graphs………………………………………………………………………. 3
**Method S2.** Pathway enrichment analysis…………………………………………………………………….... 3
**Method S3.** Spatial deconvolution analysis………………………………………………………………………. 3
**Method S4.** Intercellular communication analysis……………………………………………………………….. 4
**Method S5.** Gene network analysis……………………………………………………………………………….. 4

#### I. Supplemental Tables

**Table S1.** Overview of samples used for analyses………………………………………………………………. 5

#### II. Supplemental Figures

**Figure S1.** Histological annotation of distended and control veins………………………………………………. 6
**Figure S2.** Projection of the spatial clusters onto distended and control veins..………………………………..…. 7
**Figure S3.** Markers used for cell type identification in single-nuclei analysis……………………………………… 8
**Figure S4.** Spatial expression of cellular subpopulations in distended and control veins....…………………….. 9
**Figure S5.** RNAscope images for distended and control veins.........................................…………………….. 10
**Figure S6.** Differential intercellular interaction number and strength between distended and control veins…… 11
**Figure S7.** Immunohistochemistry staining of control, distended, and grafted veins……………………………... 12

### I. Supplemental Methods

#### Method S1. Annotation of spatial graphs

To annotate the voxels of the spatial transcriptomics plots based on histological analysis, we developed a new method using the open-access ImageJ program (v1.53t) from the National Institutes of Health. The cropped high-resolution microscopy images of the H&E stained spatial transcriptomics samples (output via Space Ranger) were imported into ImageJ. An image mask TIF was generated for each layer of the vein by selecting the outer boundary of the layer (forming a polygon) and using the Create Mask option. The three mask files for each sample were then imported into Seurat and used to add metadata labels to each spatial dataset. The function developed to add the metadata labels from the TIF images is under development and will be made available through the STannotate package via the Bhasin Lab GitHub.

#### Method S2. Pathway enrichment analysis

Pathway enrichment analyses were performed using the clusterProfiler R package (v4.6.0) and the Reactome database. The differentially expressed genes for each identity of interest, calculated using the FindAllMarkers function via Seurat, were converted to Entrez Gene IDs and used as input for the compareCluster function. The significantly enriched pathways ($P < 0.05$) were then computed for each identity using the default method of the enrichPathway function. Pathways specific to infectious disease and cancers were manually filtered.

#### Method S3. Spatial deconvolution analyses

For mapping-based integration, module scores for each single-nuclei cluster were assigned to each voxel of the spatial dataset via the ModuleScoring function in Seurat. To accomplish this, the top 15 upregulated genes for each cluster, displaying >0.5 log2 fold-change and >=30% percent cluster expression relative to the other clusters, were used as features to generate module scores (AddModuleScore function with 50 bins for scoring) for each voxel of the spatial dataset. The control and distended vein groups were then split and subset into the adventitial, medial, and intimal layers. The average module score for each subset was then calculated to generate matrices for the distended and control vein groups. A matrix of the differential module scores was then derived by subtracting the control vein matrix from the distended vein matrix.

Deconvolution analysis was also performed using Cell2location. A total of 7,518 genes were selected from the single-nuclei dataset for model training based on their expression in a minimum of five cells, expression in a minimum of 5% of the total population, and a mean non-zero expression above 1.1, using the filter_genes function. Using each sample as a variate for batch effects as well as distended versus control veins as a covariate, the annotation regression model was trained using the selected genes and 4,000 epochs through the mod.train function. The quality of the trained annotation model was then confirmed by plotting the reconstruction accuracy and mean versus estimated expression for every gene in the cluster via the mod.plot_QC function. The estimated expression for each gene for each cell population in the single-nuclei dataset (N = 10) was then extracted from the model and used for deconvolution of the spatial dataset. Genes used for the spatial mapping model (N = 6,312) were selected based on the intersection of genes between the single-nuclei and spatial datasets, wherein the latter gene set was filtered to only included genes detected in a minimum of ten spatial voxels. The spatial mapping model was then trained using each sample as a variate for batch effects, an estimated total of five cells per voxel based on the histology images, detection alpha of 200, and 10,000 epochs. The quality of the trained spatial mapping model was then confirmed by plotting the reconstruction accuracy. For each voxel, the estimated abundance for each cellular population was generated using the 5% quantile cell abundance. To generate combined spatial plots displaying the most abundant cell type for each voxel, the abundance values for each population were scaled using the MinMaxScaler function, and the corresponding cellular population with the highest scaled abundance value within each voxel was plotted.

Neighborhood enrichment analysis was then performed based on the most abundant cellular population calculated for each voxel. Using the gr.spatial_neighbors function via Squidpy, an enrichment score was generated for each cellular population based on their spatial proximity, where a high score represents closer proximity. To build the enrichment plot, a connectivity matrix was then generated using the gr.nhood_enrichment function and plotted via the pl.nhood_enrichment function.

**Method S4. Intercellular communication analysis**

Intercellular communication analysis was performed using CellChat (v1.6.1).13 The single-nuclei dataset was subset into the distended vein and control vein groups, which were analyzed first as separate CellChat objects following previously described methods.14 The objects were then combined via mergeCellChat to compare the differences in their intercellular communication networks. The differential heatmap of interactions was generated by subtracting the overall interaction matrix of the control veins from the distended veins.

**Method S5. Gene network analysis**

Gene network analysis was performed using Cytoscape (v3.10.0). Differentially expressed genes (adj. p-value $< 0.05$, average $\log_2FC > 0$) for each group (control veins, distended veins, and grafted veins) were calculated using the FindMarkers function via Seurat. These genes were then used as input nodes to construct individual STRING networks for each group, with 13 nodes and 0 edges for the control gene network, 100 nodes and 175 edges for the distended gene network, and 851 nodes and 9,650 edges for the grafted gene network. To connect these networks as layers within a temporal network, interlayer edges connecting the nodes of the control and distended network layers were calculated using STRING, in addition to the interlayer edges connecting the nodes of the distended and grafted network layers. TimeNexus was then used to construct a multilayer network based on these inputs with a total of 964 nodes and 10,496 edges. To filter the network for targeted analysis, the nearest intralayer and interlayer neighbors of FBN1, GLIS3, and VCAN were calculated using the analyze network function and the remaining nodes and edges were hidden.

### II. Supplemental Tables

**Table S1. Overview of samples used for analyses.**

| | Canine 1 | Canine 2 | Canine 3 | Canine 4 |
|---|---|---|---|---|
| **Control vein samples:** | | | | |
| Single-nuclei RNA-seq sample name *(no. nuclei captured)* | C1_SN1 *(609 nuclei)* | C2_SN1 *(579 nuclei)* | – | – |
| Spatial transcriptomics sample name *(no. spots captured)* | C1_ST1 *(330 spots)* | C2_ST1 *(355 spots)* | C3_ST1 *(157 spots)* | C4_ST1 *(127 spots)* |
| **Distended vein samples:** | | | | |
| Single-nuclei RNA-seq sample name *(no. nuclei captured)* | C1_SN2 *(1000 nuclei)* | C2_SN2 *(223 nuclei)* | – | – |
| Spatial transcriptomics sample name *(no. spots captured)* | – | C2_ST2 *(541 spots)* | C3_ST2 *(353 spots)* | C4_ST2 *(321 spots)* |

### III. Supplemental Figures

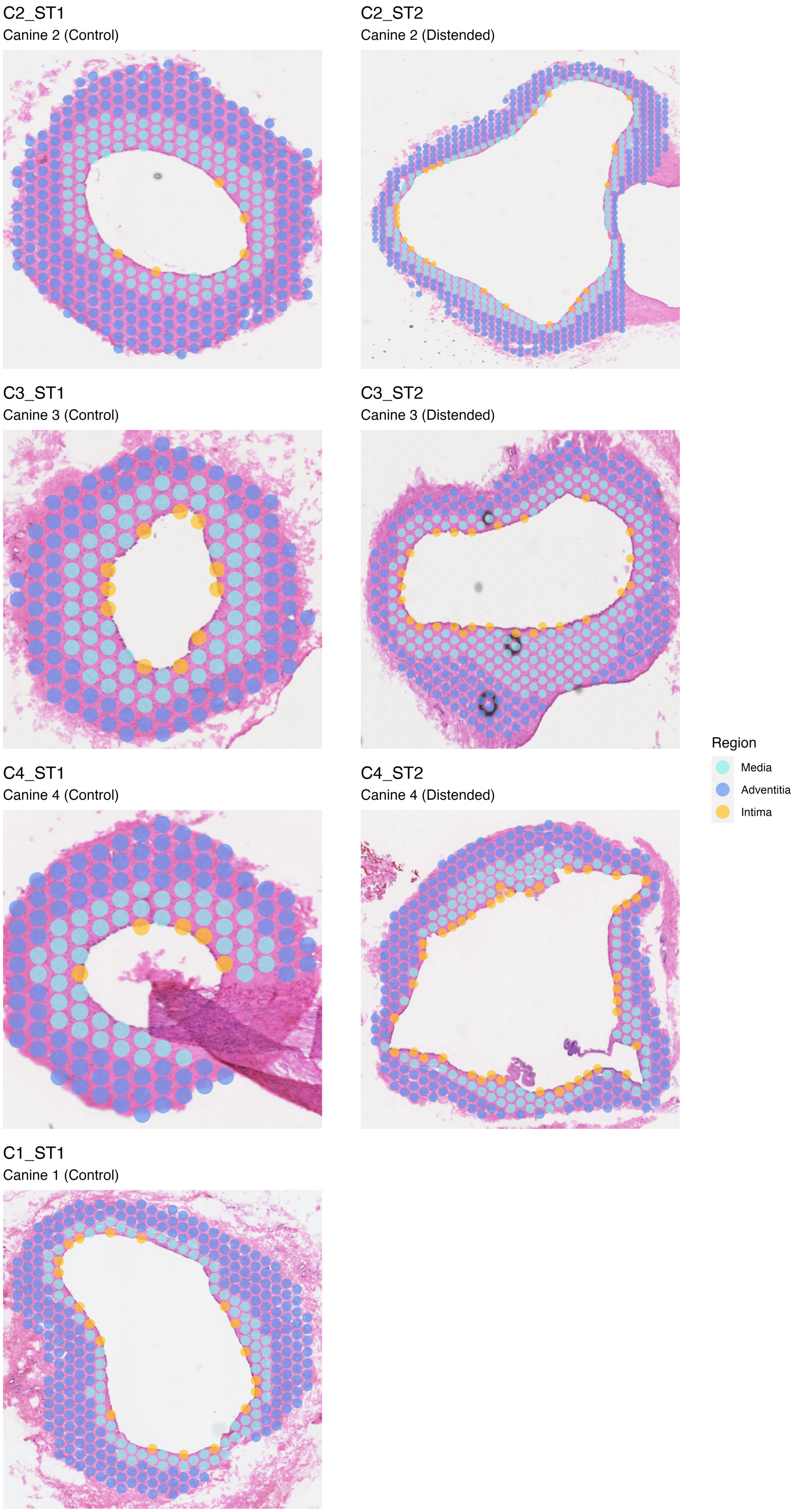


**Figure S1. Histological annotation of distended and control veins.** The layers of the vein wall were annotated for both control and distended vein samples using the developed histological annotation tool.

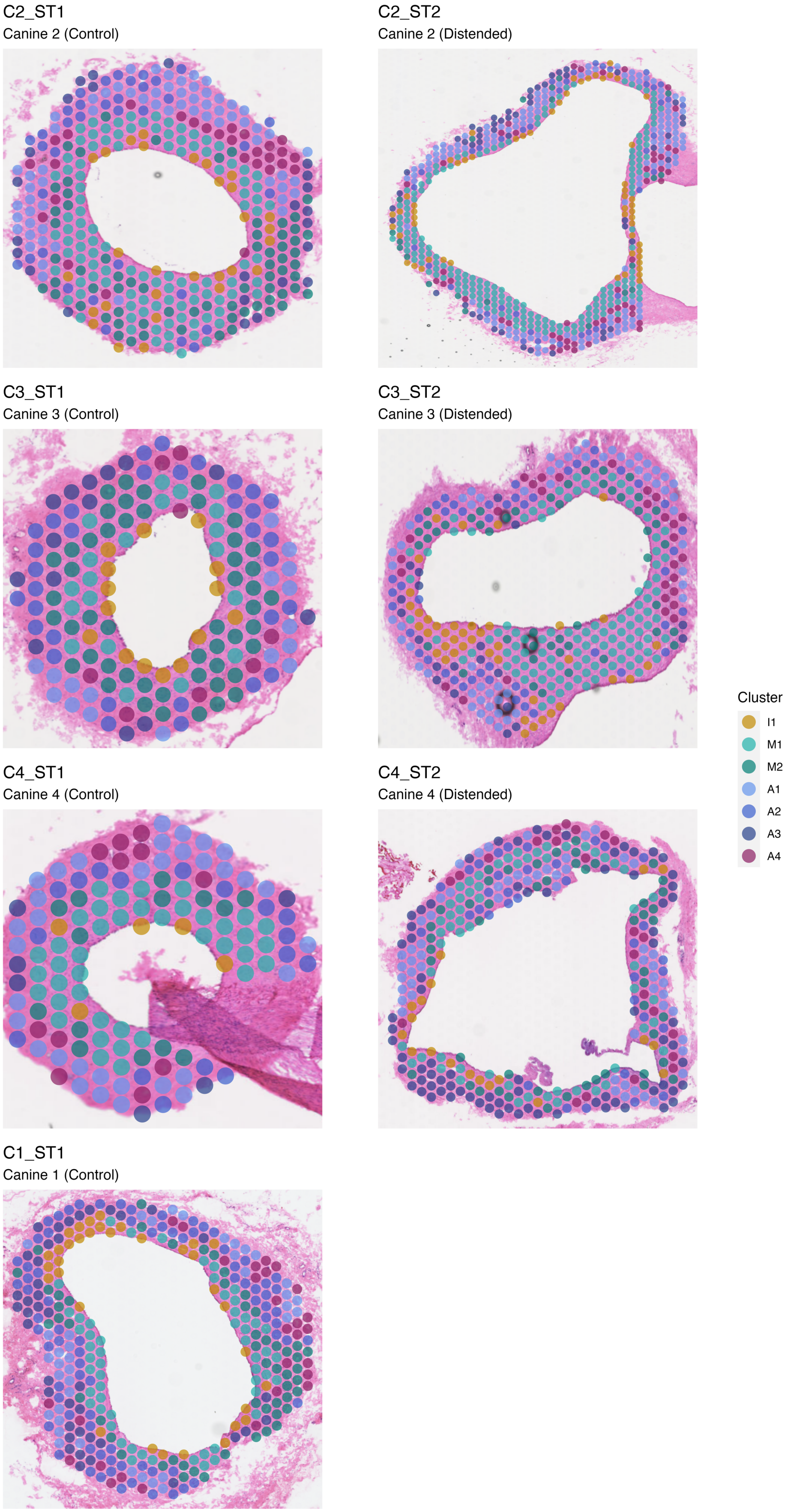


**Figure S2. Projection of the spatial clusters onto distended and control veins.** The layers of the vein wall were annotated for both control and distended vein samples using the developed histological annotation tool. Different spatial clusters from Intima, Media, and Adventitia were projected on spatial vein tissues to learn about spatial distribution.

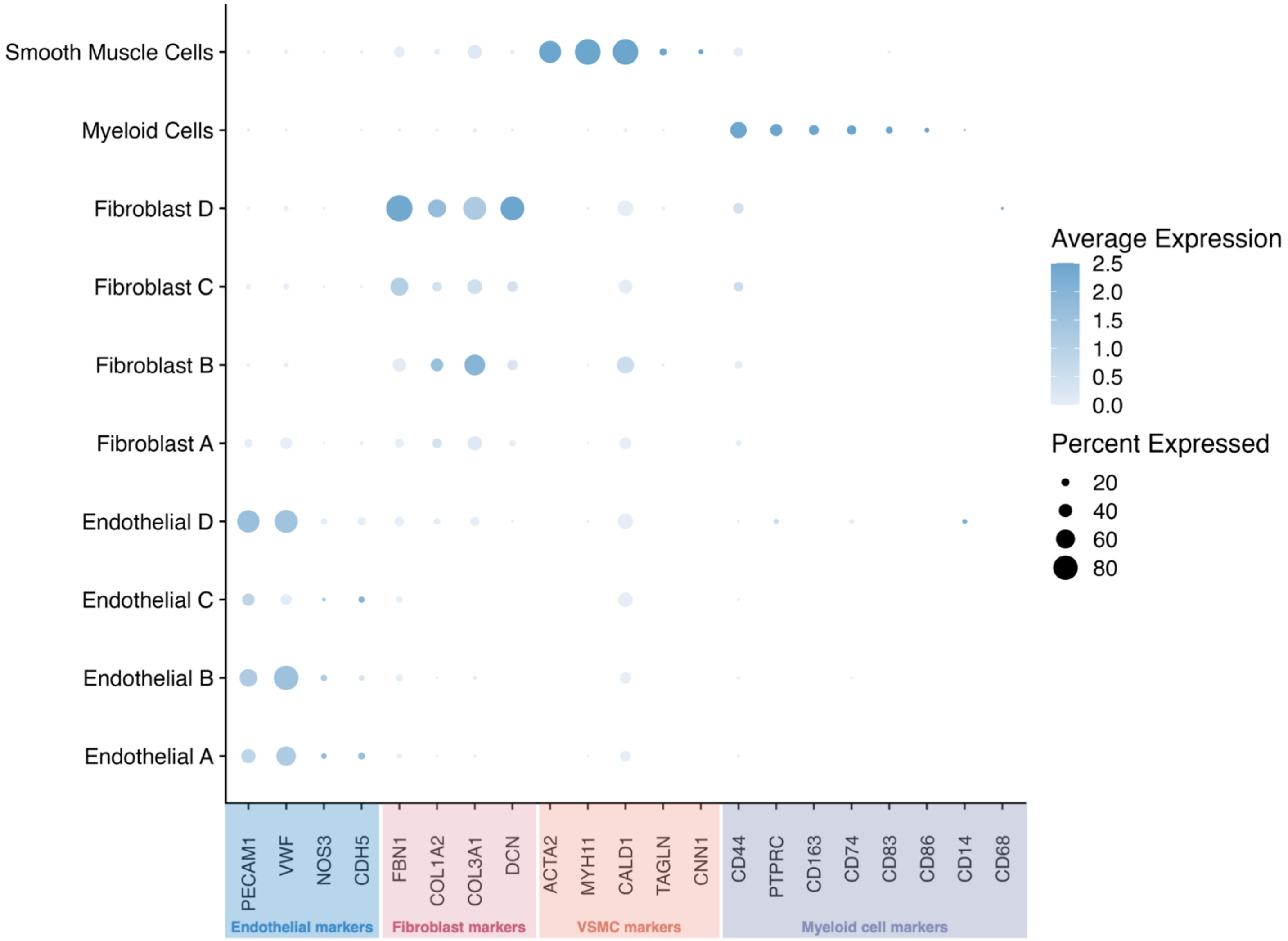


**Figure S3. Markers used for cell type identification in single-nuclei analysis.** Clusters identified through single-nuclei analysis were labeled as either endothelial, fibroblast, VSMC, or myeloid cells, based on their expression of canonical markers associated with these cell types.

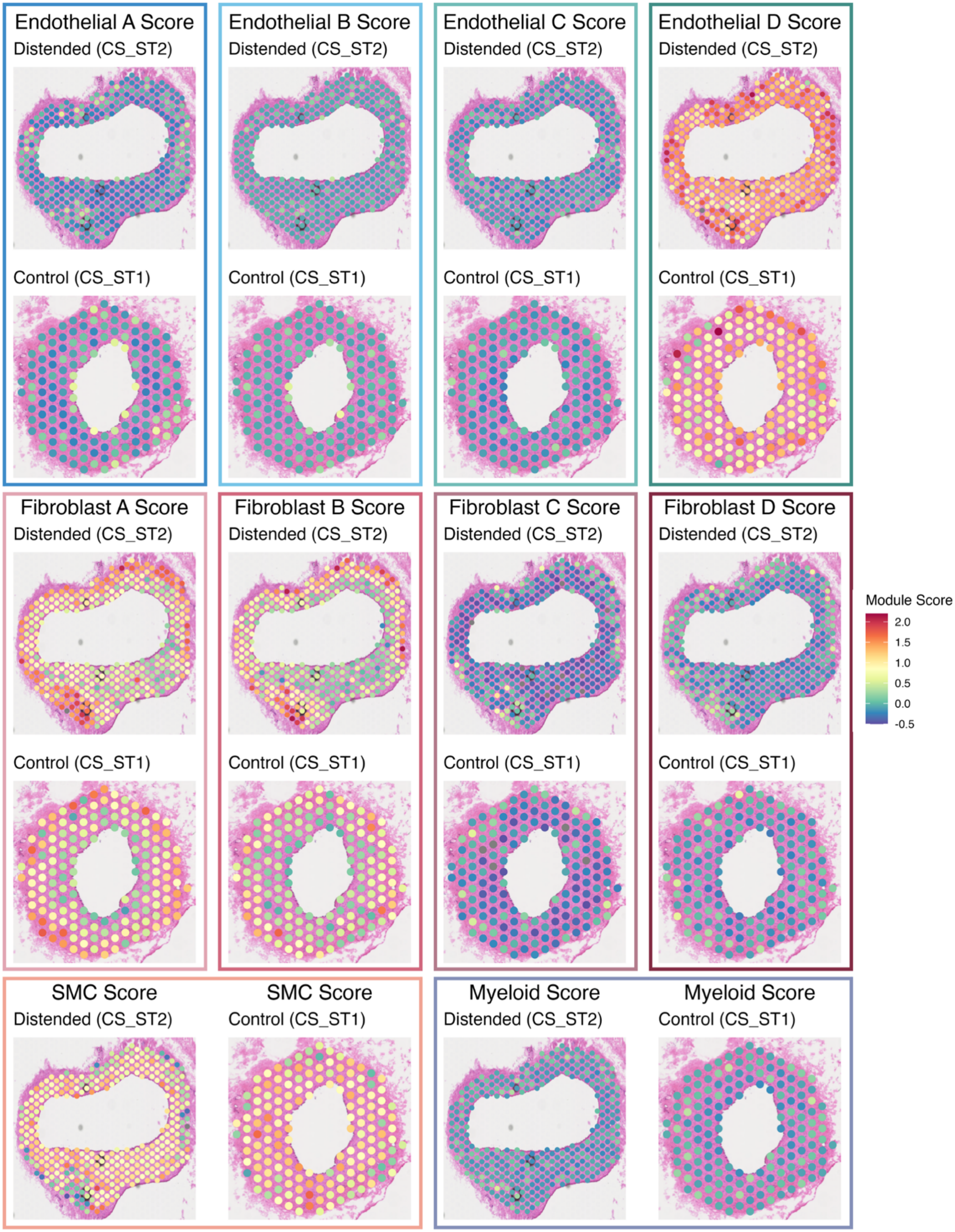


**Figure S4. Spatial expression of cellular subpopulations in distended and control veins.** Spatial plots displaying the module scores for each cellular subpopulation identified through snRNA-seq in paired control and distended vein samples.

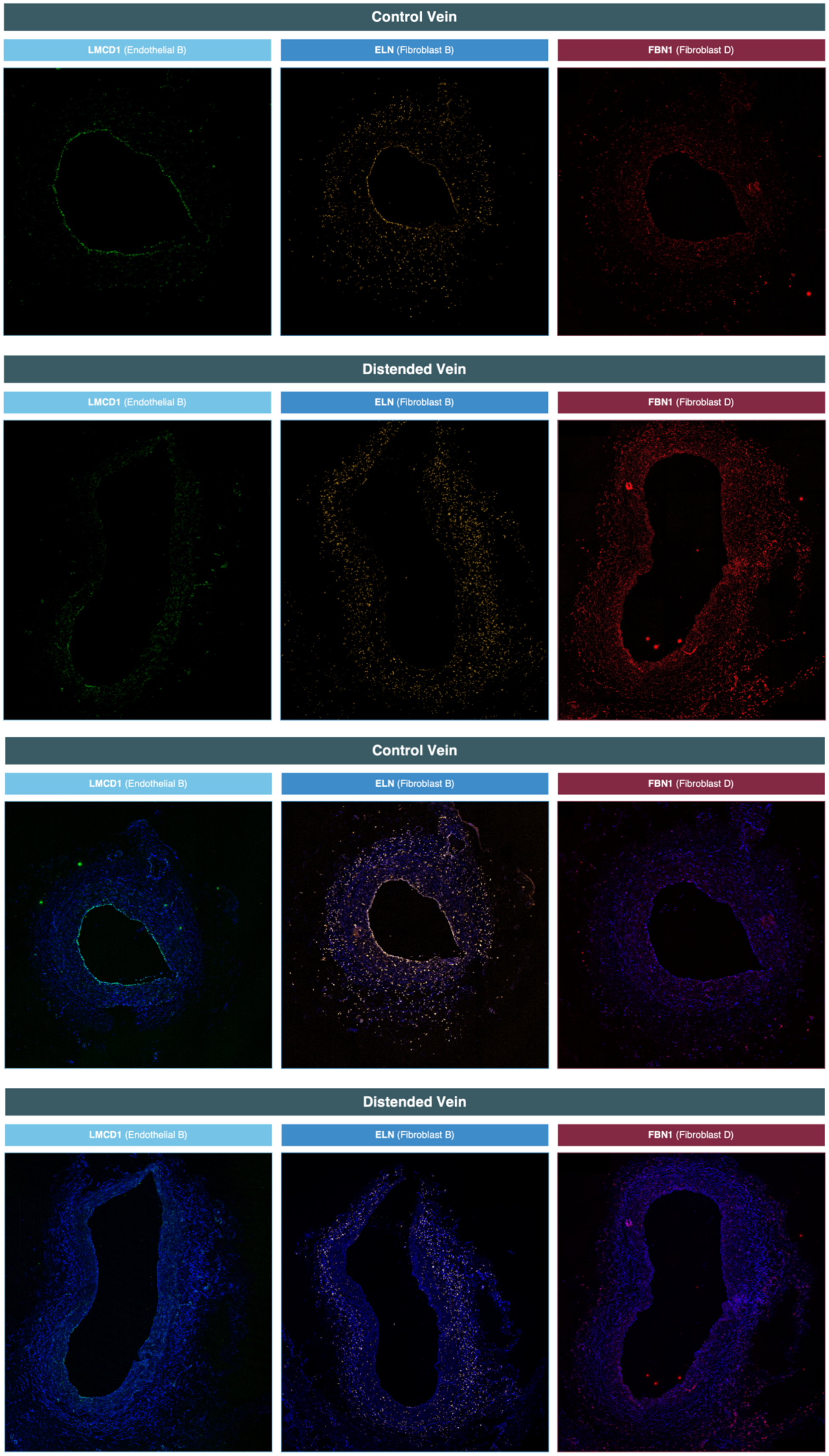


**Figure S5.** Confocal images of LMCD1 (green), ELN (yellow), and FBN1 (red) RNAscope probe fluorescence in isolation (top) and merged with DAPI (bottom). Probes were selected as representative markers of the Endothelial B, Fibroblast B, and Endothelial D subpopulations elucidated via single-nuclei analysis. Elastin is captured in the intima of control veins but not distended veins, likely due to the loss of intimal ECs following distension.

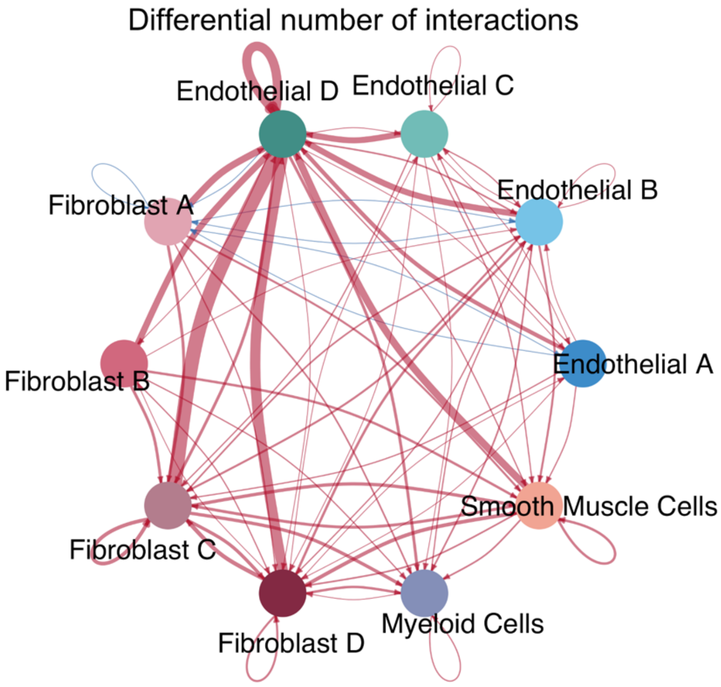


**Figure S6. Differential intercellular interaction number and strength between distended and control veins.** Differential number of intercellular communication interactions between subpopulations, wherein red and blue indicate an increased or decreased, respectively, number of interactions within distended veins compared to control veins. Differential strength of the intercellular communication interactions between subpopulations, wherein red and blue indicate an increased or decreased, respectively, strength of signaling interactions between subpopulations in distended veins compared to control veins.

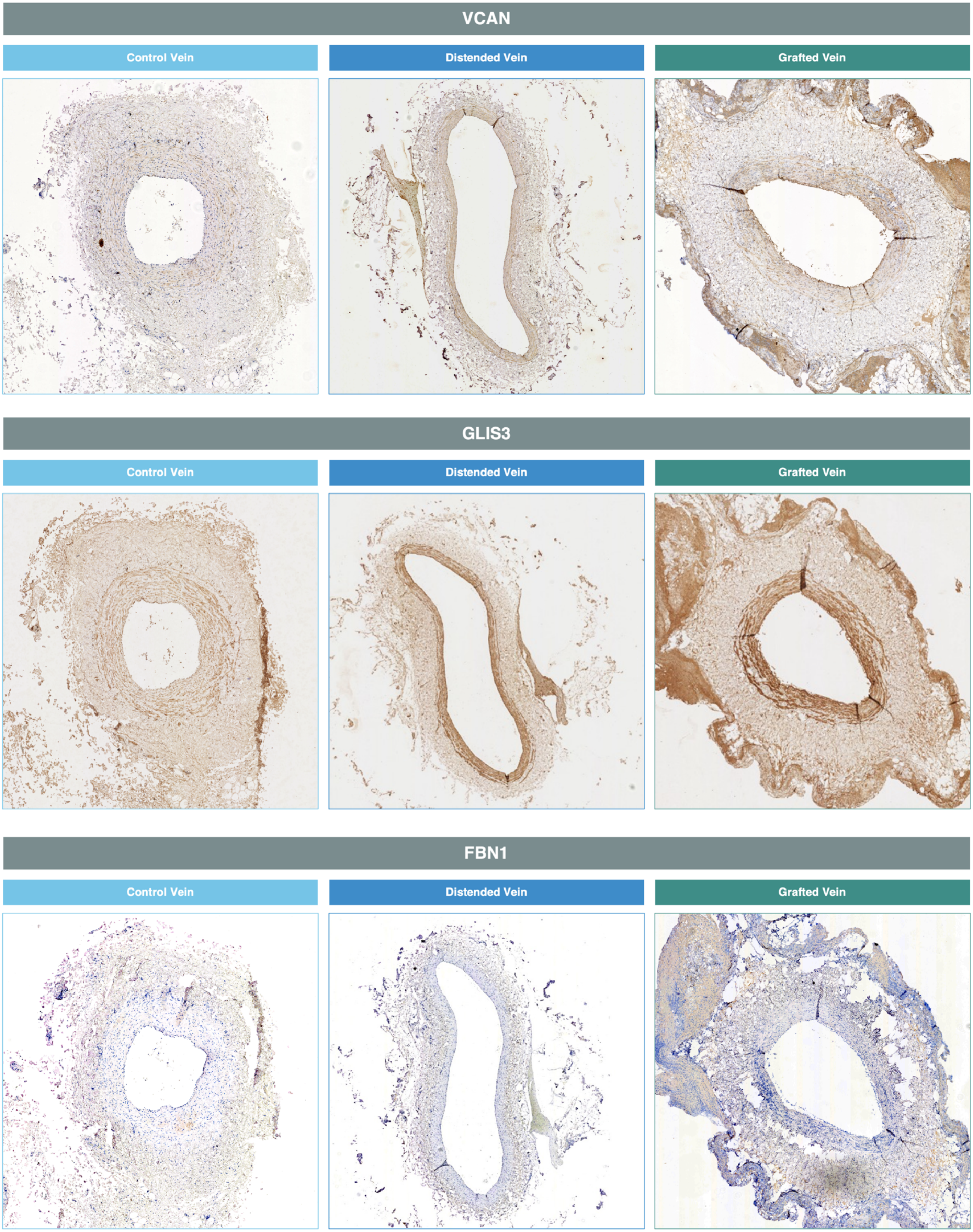


**Figure S7.** Immunohistochemistry staining of control, distended, and grafted veins.
